# Morpheus-3D: Structural Diversity-Guided Detection and Localization of Protein Fold Switching

**DOI:** 10.64898/2026.08.16.745091

**Authors:** Sreeharsh Kuniyil, Vijay Subramanian, Akanksha Arun, Anand Lakshmanan, Ashok Sekhar, Anand Srivastava

## Abstract

Proteins that reversibly adopt multiple stable folds challenge the classical sequence–structure paradigm, yet their discovery remains limited because fold switching is difficult to detect experimentally and current computational methods fail to resolve the underlying conformationally plastic regions. Here we present Morpheus-3D, a sequence-based framework that quantifies residue-level tertiary structural diversity using entropy profiles derived from the Foldseek 3Di structural alphabet. By capturing variation in tertiary interaction environments rather than secondary structure alone, Morpheus-3D identifies fold-switching proteins while simultaneously localizing the sequence regions responsible for structural transitions. The framework outperforms existing predictors, accurately recovers experimentally characterized switching regions, generalizes to recently discovered natural and engineered fold-switching proteins absent from training, and detects conformational plasticity inaccessible to secondary-structure-based approaches. Application to 57 representative proteomes reveals that fold-switching potential is widespread but enriched in regulatory, pathogenic, and environmentally adaptive lineages. Integration with ancestral sequence reconstruction further uncovers evolutionary trajectories through which conformational plasticity emerges. To make these predictions directly accessible, we implemented Morpheus-3D as an interactive web platform (https://morpheus.slicearrow.com/), in which per-residue entropy profiles, sequence and three-dimensional structure are displayed together and respond as one, allowing predicted fold-switching regions to be mapped onto the structure and exported for downstream analysis. Morpheus-3D provides a scalable framework for discovering metamorphic proteins and investigating the origins, mechanisms, and evolution of structural plasticity directly from sequence.

## Introduction

The classical sequence-structure-function paradigm, established by Anfinsen, holds that a protein’s amino acid sequence uniquely determines its three-dimensional structure and thereby its biological function (1). Most globular proteins conform to this principle, but several classes deviate substantially. Metamorphic proteins represent a distinct category in which a single amino acid sequence reversibly adopts two or more thermodynamically stable folded conformations that differ in secondary or tertiary structural organization and are frequently associated with distinct biological functions (2, 3). Unlike intrinsically disordered proteins (IDPs), which lack stable folded states, metamorphic proteins interconvert between distinct, well-folded conformations and unlike irreversible fold switchers, this interconversion is fully reversible (4, 5).

Experimentally characterized metamorphic proteins span diverse biological contexts and illustrate the functional importance of conformational plasticity. The bacterial transcription factor RfaH and the cyanobacterial circadian regulator KaiB are among the most extensively studied examples, each reversibly switching between two folded conformations to control a distinct biological function (3, 6, 7). The human chemokine XCL1 interconverts between a canonical chemokine fold that activates XCR1 and an alternative all-β dimer that binds glycosaminoglycans, partitioning distinct signalling functions between two folded states (8, 9). Conformational switching in these and related proteins is regulated by stimuli including temperature, ionic strength, ligand binding and redox state, establishing structural plasticity as a conserved and functionally consequential regulatory mechanism. Despite this growing body of evidence, fewer than one hundred metamorphic proteins have been conclusively characterized. Prior estimates suggest that metamorphic proteins may constitute between 0.5 and 4 percent of experimentally determined protein structures, implying that the known repertoire captures only a small fraction of the complete metamorphome (10). Experimental identification of alternative conformations requires nuclear magnetic resonance spectroscopy, X-ray crystallography under varied experimental conditions, or cryogenic electron microscopy, each inherently low-throughput and unsuitable for proteome-scale discovery. Computational approaches are therefore essential for systematic candidate identification and estimation of metamorphic protein prevalence across complete proteomes.

Several sequence- and structure-based computational strategies have been developed toward this goal. Chen et al. introduced a diversity index quantifying disagreement among predicted secondary structure probabilities, demonstrating that elevated structural ambiguity is associated with fold-switching propensity (11). This approach established that sequence-derived structural uncertainty carries predictive information, but is limited to secondary structure signals and cannot detect conformational rearrangements that alter tertiary interaction networks while preserving local backbone geometry. Methods addressing specific structural subclasses, including α-helix to β-strand conversion (12) and sequence-similar fold switchers arising through evolutionary divergence (13), cover narrow transition types or address homologous sequence variation rather than reversible interconversion within a single sequence. AlphaFold-based conformational sampling through manipulation of multiple sequence alignments or model inference parameters has expanded the structural hypotheses available for fold-switching candidates (14–16), but requires repeated structure prediction and incurs computational costs that preclude routine proteome-scale application. Whether recovered alternative conformations reflect genuine model generalization or memorization of deposited experimental structures also remains unresolved (17–19).

Complementary to sequence-based prediction, molecular dynamics simulation approaches have been implemented to characterize the thermodynamic and kinetic basis of metamorphic interconversion directly. Enhanced-sampling molecular dynamics methods, including hybrid parallel tempering schemes, have enabled high-resolution ensemble reconstruction of the metamorphic protein RfaH recovering multi-funneled free energy landscapes consistent with NMR measurements (20, 21). Free energy calculations combining advanced sampling with explicit solvent thermodynamics have further resolved the enthalpic and entropic driving forces underlying temperature-induced fold switching in the designed metamorphic protein Sa1_V90T (22), showing that release of ordered water governs the transition between alternative folds (23). At a more general level, statistical mechanical folding models incorporating multiple native basins have unified metamorphicity, intrinsic disorder and folding-upon-binding within a common theoretical framework (24). Together, these studies provide detailed mechanistic and thermodynamic insight into individual fold-switching systems but remain computationally intensive on a per-protein basis and are not designed for proteome-scale screening.

Our earlier framework, Morpheus-1, addressed proteome-scale screening by computing residue-level secondary structure entropy over sequence fragments, quantifying the degree to which a given fragment occupies multiple secondary structural contexts across experimental and predicted protein structures (25). However, secondary structure entropy is insensitive to conformational rearrangements that remodel tertiary interaction networks without altering local backbone geometry, reducing sensitivity for an important subset of metamorphic proteins. Furthermore, neither Morpheus-1 nor existing methods provide residue-level localization of fold-switching regions, limiting their utility for mechanistic interpretation and experimental design. The Foldseek three-dimensional interaction (3Di) structural alphabet addresses this gap by assigning each residue to one of twenty discrete states defined by the spatial geometry of interactions with its nearest structural neighbour, encoding tertiary contact information absent from conventional secondary structure representations (26). Entropy computed within the 3Di representation reflects variation in tertiary interaction environments and captures conformational rearrangements invisible to secondary structure analysis. These two descriptors are complementary and together provide a substantially more complete characterization of structural heterogeneity than either representation alone.

Here we present Morpheus-3D, a fragment-based framework that integrates 3Di tertiary interaction diversity with secondary structure entropy for sequence-based prediction of fold-switching proteins and residue-level localization of switching regions. The overall workflow of the Morpheus-3D pipeline is illustrated in Fig. 1. Morpheus-3D out-performs existing sequence-based prediction methods, with the largest gains for proteins that undergo tertiary rearrangement while preserving secondary structure composition. Application to 57 representative proteomes spanning six kingdoms characterizes the distribution of conformational plasticity across the tree of life. To investigate the evolutionary origins of fold switching, we further integrated ancestral sequence reconstruction with residue-level entropy profiling, demonstrating that the emergence of conformational plasticity can be traced through phylogenetic lineages and associated with specific mutational events along ancestral branches. All source code, trained models and curated databases are made publicly available.

## Methodology

### 3D Structure Alphabet Representation

To enable scalable structural comparison across large protein databases, we encode three-dimensional protein structures as discrete sequences using the Foldseek 3Di structural alphabet (26). This encoding converts continuous structural space into a symbolic representation amenable to sequence-based alignment algorithms, dramatically reducing the computational cost of structure search without proportional loss in sensitivity. The 3Di alphabet comprises 20 discrete states, each describing the geometric relationship between a residue *i* and its nearest spatial neighbor *j*, identified via a learned virtual center position defined by angle *θ*, dihedral angle *τ*, and distance *l* from the C*α* atom. Crucially, this nearest-neighbour criterion prioritizes long-range tertiary contacts over local backbone neighbours. For each residue pair (*i, j*), the geometric conformation is captured through a 10-dimensional feature vector comprising seven inter-backbone angular cosines derived from unit vectors along the C*α* backbone fragments of both residues, the Euclidean C*α*–C*α* distance, and two sequence distance features encoding the relative positions of *i* and *j* in the primary sequence. These features are discretised into one of 20 states using a Vector Quantized Variational Autoencoder (VQ-VAE) (27), trained to maximise evolutionary conservation of the learned states, with training performed on structurally aligned residue pairs from the SCOPe40 dataset (28). In this work, the 3Di encoding serves as the primary structural abstraction for all downstream indexing, retrieval, and similarity computation, enabling database-scale queries that would otherwise be computationally intractable.

**Fig. 1.**
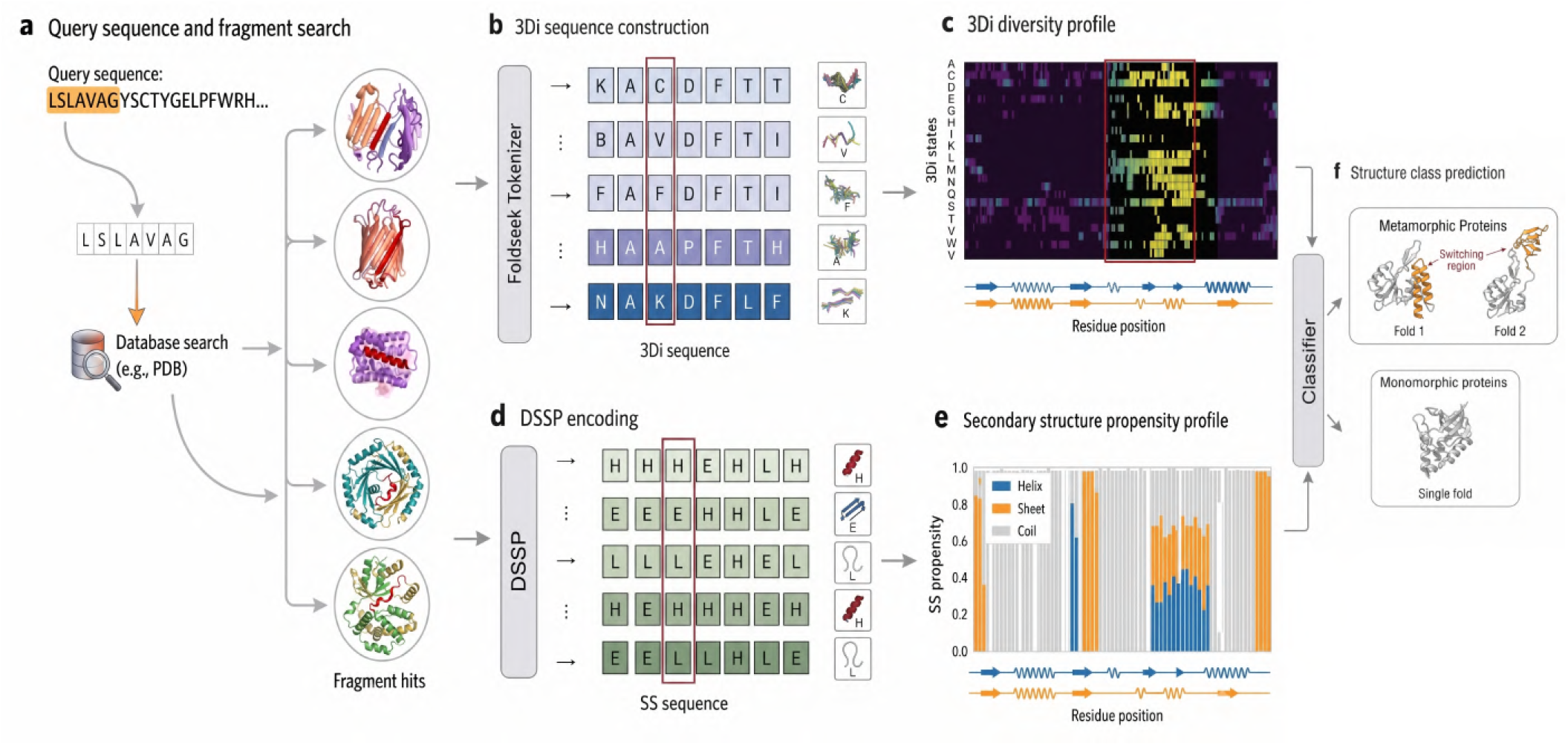
Overview of the Morpheus-3D framework. (**a**) The query protein sequence is divided into overlapping 7-mer fragments and searched against a structure-informed database containing experimentally determined and AlphaFold-predicted protein structures. (**b**) Retrieved fragments are encoded using the Foldseek 3Di structural alphabet to generate residue-level tertiary interaction states. (**c**) Shannon entropy of the 3Di state distribution is computed to produce a residue-level tertiary structural diversity profile. (**d**) Secondary structure annotations are obtained using DSSP. (**e**) Secondary structure propensities are calculated to generate residue-level profiles of helix, strand and loop preferences. (**f**) Features derived from the 3Di diversity and secondary structure profiles are integrated by a supervised machine-learning classifier to identify metamorphic proteins and localize putative fold-switching regions.

### Database Construction

We constructed a comprehensive structure-informed protein database by integrating experimentally resolved and computationally predicted protein structures. A total of 200,000 redundant protein structures were obtained from the Protein Data Bank (29), along with 1.2 million structure predictions generated by AlphaFold2 (30). All structures were processed using Foldseek to derive 3Di structural alphabet sequences. This enables efficient indexing, comparison, and search using sequence-based methods. To enrich the structural representation, secondary structure annotations were assigned using DSSP (31), generating residue-level classifications. For AlphaFold-predicted structures, per-residue confidence scores (pLDDT) were additionally extracted to quantify prediction reliability. These features were systematically integrated to produce a unified, residue-aligned dataset comprising: (i) primary amino acid sequences, (ii) Foldseek-derived 3Di structural sequences, (iii) DSSP-based secondary structure annotations, and (iv) confidence scores for predicted structures. The resulting database forms a multi-modal representation of protein structure and sequence, designed to support downstream tasks. The curated dataset is made publicly available to facilitate reproducibility and further research.

#### Training Data Curation

The training dataset comprises 94 experimentally verified metamorphic proteins as the positive class and 94 monomorphic proteins confirmed to adopt a single stable fold, drawn from established curated benchmarks (10, 11) as the negative class, for a combined dataset of 188 proteins. The two classes are approximately balanced, supporting unbiased classifier training. A fundamental challenge in this domain is the extreme scarcity of confirmed metamorphic proteins, fewer than a hundred experimentally validated examples exist. High-confidence positive predictions on unseen sequences should therefore be interpreted as candidates for experimental follow-up rather than definitive classifications. Despite this constraint, the dataset captures meaningful structural diversity. The positive set spans a broad sequence length range (26–800 residues) and encompasses diverse fold-switching mechanisms, including ligand-induced conformational switching, and environment-driven structural transitions such as pH or redox dependent refolding, ensuring the classifier is not biased toward any single mode of conformational change.

#### Fragment-Based Structural Retrieval

The query sequence was segmented into overlapping 7-mer fragments with a stride of one residue, each queried against the curated database for identical sequence matches. The database stores, for each entry, the amino acid sequence, Foldseek-derived 3Di sequence, secondary structure annotation and per-residue confidence scores. Tertiary interaction context and secondary structure propensity for each retrieved hit were obtained from the corresponding 3Di and secondary structure 7-mers respectively. The fragment length k = 7 was determined empirically by balanced one-way ANOVA (32) performed across fragment sizes ranging from 5 to 8 residues using the maximum rolling 3Di entropy as the discriminating feature. To eliminate class imbalance, monomorphic proteins were randomly subsampled to match the number of metamorphic proteins, and the analysis was repeated across 100 bootstrap replicates. Seven-residue fragments consistently produced the highest mean F-statistics and were therefore adopted for all subsequent analyses (Supplementary Fig. S1). To support efficient proteome-scale retrieval, all unique 7-mers in the database were indexed using a SQLite-based lookup table (33) mapping each fragment to its associated protein identifiers, 3Di sequences, secondary structure sequences and per-residue confidence scores. Retrieved hits were filtered to exclude fragments in which the central residue corresponded to an unobserved 3Di or secondary structure state or exhibited low structural confidence (pLDDT < 70 for predicted structures). Additionally, to ensure that fragment-level statistics are not biased by over-represented sequences, the database was clustered at 100% sequence identity using MMseqs2 (34). The resulting filtered, non-redundant fragment hits were subsequently used for downstream scoring and diversity analysis.

#### Scoring Metrics

For each retrieved hit, the 3Di state and secondary structure assignment are recorded at the 3rd residue of the fragment. The representative residue position within each 7-mer was selected using the same balanced one-way ANOVA (32) framework. Among all candidate positions, the third residue consistently produced the highest mean F-statistic across 100 balanced bootstrap replicates and was therefore selected for all subsequent analyses (Supplementary Fig. S1). The central novelty of Morpheus-3D is the introduction of a tertiary interaction diversity score derived from the Foldseek 3Di structural alphabet. When the same sequence fragment maps to hits spanning multiple distinct 3Di states, it reflects tolerance to multiple tertiary interaction environments, a hallmark of conformationally plastic regions; a narrow distribution conversely indicates a structurally rigid locus. This diversity is quantified by treating the 3Di state distribution across all retrieved hits as a discrete probability distribution over the 20 possible states and computing its Shannon entropy (35):

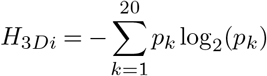

where *p*_*k*_ is the fractional occupancy of the *k*-th 3Di state. A high *H*_3*Di*_ value indicates structural ambiguity in the tertiary interaction context of that fragment across the database, while a low value reflects a consistent and rigid tertiary environment. Critically, *H*_3*Di*_ captures conformational rear-rangements that alter tertiary interaction geometry without remodeling local backbone conformation, a class of transitions to which secondary structure metrics are insensitive. Secondary structure diversity is assessed from the fractional propensities of helix (*h*), strand (*e*) and loop (*l*) states across all retrieved hits. Two complementary metrics are computed: the diversity index (DI), originally proposed by Chen et al. (11), and the secondary structure entropy (*H*_*SS*_), introduced in Morpheus-1 (25):

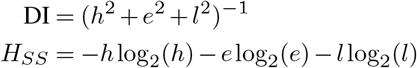

Higher values of both metrics indicate greater secondary structural ambiguity at that position. All three metrics were summarised over rolling windows ranging from 5 to 20 residues. Balanced one-way ANOVA (32) demonstrated that larger windows progressively improved discrimination between metamorphic and monomorphic proteins, with a window size of 20 residues yielding the highest mean F-statistic. This value was therefore used throughout the study (Supplementary Fig. S1). For each metric, both the rolling maximum and rolling mean are computed. The rolling maximum captures the peak structural diversity within the sequence, reflecting the most conformationally ambiguous local region. The rolling mean carries richer information about the extent and distribution of structural heterogeneity across the sequence, better representing sustained stretches of elevated entropy that are characteristic of fold-switching regions. The resulting six features form a unified multi-resolution representation jointly characterising tertiary interaction diversity and secondary structure heterogeneity, constituting the input feature space for downstream classification.

#### Model Development and Evaluation

The feature vector for each protein comprises six scores derived from the fragment scoring pipeline: the rolling mean and maximum of the 3Di-derived tertiary interaction entropy (*H*_3*Di*_), and the rolling mean and maximum of the secondary structure diversity index (DI) and entropy (*H*_*SS*_). To identify the most suitable classification model, four canonical machine learning algorithms were evaluated: Logistic Regression (36), Support Vector Machine (SVM) (37), Random Forest (38) and Gradient Boosting (39). This comparison ensures that model selection is not biased toward any single algorithmic family and reflects a principled, evidence-based choice. Model training and evaluation were performed using a nested cross-validation scheme comprising a 6-fold outer loop and a 3-fold inner loop (Supplementary Fig. S2). The outer loop provides an unbiased estimate of generalisation performance on data entirely withheld during training, while the inner loop is used exclusively for hyperparameter optimisation. This separation ensures that model selection and performance estimation remain independent, preventing information leakage between tuning and evaluation. Performance across outer folds was assessed using accuracy, precision, recall, F1-score and Matthews Correlation Coefficient (MCC). Among all models evaluated, SVM with an RBF kernel consistently achieved the highest and most stable performance across outer folds, exhibiting the most favourable balance between training and held-out accuracy, minimal overfitting and strong discriminative performance across all evaluation metrics (Supplementary Fig. S3). It was accordingly selected as the final classification model.

#### Proteome-wide Screening and Functional Annotation

Proteome FASTA sequences from 57 representative species spanning six major taxonomic groups (Eubacteria, Archaea, Animalia, Plantae, Fungi, and Protists), together with a representative viral proteome, were downloaded from the UniProt Proteomes database (40). Each proteome was analysed using the Morpheus-3D pipeline to generate residue-level 3Di and secondary-structure entropy profiles. A residue was classified as a putative fold-switching position if a 20-residue centered rolling average of its raw *H*_3*Di*_ and *H*_*SS*_ values (in bits) both exceeded empirically chosen cutoffs. These cutoffs were determined using a benchmark dataset of experimentally annotated fold-switching residues within known fold-switching proteins. Candidate threshold values were scanned across the observed entropy range, and the resulting false-positive and false-negative residue counts were quantified at each value (Supplementary Fig. S4), revealing the expected trade-off between the two error types. Thresholds of 0.95 bits for *H*_3*Di*_ and 0.40 bits for *H*_*SS*_ were selected as recall-oriented operating points, prioritising recovery of true fold-switching residues while keeping the false-positive rate acceptable. Hotspot regions from the two rolling-averaged tracks were reconciled into a single consensus fold-switching region by intersecting and merging their overlapping intervals. All predicted metamorphic proteins were subsequently analysed using IUPred2A (41) and MetaPredict (42). Proteins in which more than 50% of residues within the predicted switching region were consistently classified as intrinsically disordered by both methods were excluded from further analyses. Functional annotation was performed using Gene Ontology (GO) Biological Process annotations obtained from UniProt (40). GO terms were mapped to organism-specific high-level biological process categories using the Gene Ontology hierarchy implemented in GOATOOLS (43), and proteins assigned to multiple categories were weighted by fractional counting before category frequencies were summarised for each proteome.

### Evolutionary Analysis Using Ancestral Sequence Reconstruction

Homologous sequences were retrieved from InterPro (44) for three protein families selected for evolutionary analysis. For XCL1, 11,977 sequences belonging to the Beta/Gamma/Delta Chemokine family (InterPro: IPR039809) were collected, spanning both XCL1 and CCL20 lineages. For KaiB, 2,194 sequences were obtained from the Circadian Clock Protein KaiB-like family (InterPro: IPR039022). For RfaH, 31,206 sequences were retrieved from the NusG-like family (InterPro: IPR043425). Redundant entries were removed by clustering each dataset at 100% sequence identity using CD-HIT (45) prior to phylogenetic analysis. Multiple sequence alignments were generated using MAFFT (46) and subsequently refined with ClipKIT (47) using a gap threshold of 0.7 to remove poorly aligned positions while retaining phylogenetically informative sites. Maximum-likelihood phylogenetic trees were constructed using IQ-TREE (48) under the LG+G substitution model (49). Ancestral sequences at internal phylogenetic nodes were reconstructed using the maximum-likelihood ancestral state reconstruction framework implemented in IQ-TREE. Reconstructed ancestral sequences were subsequently passed through the Morpheus-3D pipeline to generate residue-level structural diversity profiles, enabling systematic evaluation of evolutionary changes in conformational plasticity across phylogenetic lineages.

### The Morpheus-3D Web Platform

To make residue-level predictions directly explorable, the Morpheus-3D pipeline was implemented as an interactive web platform (https://morpheus.slicearrow.com/). An analysis begins from a protein sequence, supplied either as direct FASTA input or fetched by accession from the PDB (29) or UniProt (40) databases, and returns the predicted fold-switching class, summary features, and an estimated probability of multi-fold behaviour. Per-residue results appear across three linked panels: a sequence feature graph plotting rolling *H*_3*Di*_ and *H*_*SS*_ profiles, from which a secondary-structure diversity plot and a per-position 3Di state probability heatmap open; a sequence viewer in register with these profiles; and a structure viewer built on the NGL engine (50) with user-defined representations, colour schemes and projection. Hovering or clicking a residue highlights it across all panels and collates selections into a per-residue entropy table. For each query, an AlphaFold2 model is retrieved from the AlphaFold Protein Structure Database and aligned to the submitted sequence by global pairwise alignment using Biopython (51), classified as an exact, partial, similar, or absent match; where no match is found, a model is generated on demand with ESM-Fold (52), returning a per-residue pLDDT profile. Entropy hotspots and fold-switching regions can be rendered onto the structure on a blue–white–red scale, with multi-chain assemblies mapped via per-chain BLOSUM62 alignment. For PDB-derived structures, deposited coordinates and the predicted model are displayed interchangeably, with experimental method and resolution retrieved from the RCSB Data API (53), and bound ligands, ions and waters shown or hidden individually. Rendered views export as PNG, and complete pipeline outputs download as a compressed archive.

## Results

### A. Incorporating tertiary structure diversity enhances fold-switching prediction accuracy

To evaluate the contribution of tertiary structural information to fold-switching prediction, Morpheus-3D was benchmarked against the existing sequence-based classifier of Chen et al. (11) and its direct predecessor, Morpheus-1 (25). Both prior approaches treat secondary structure diversity as the primary basis for predicting conformational plasticity. Morpheus-3D builds upon this foundation by introducing 3Di-derived tertiary interaction diversity as an additional predictive layer, directly testing whether structural information beyond secondary organisation carries independent discriminative value. Morpheus-3D demonstrated consistent improvements across all evaluation metrics relative to both baselines (Fig. 2a). Compared to Chen et al., gains were substantial, with MCC more than doubling. Against Morpheus-1 (25), accuracy improved by approximately 2.4% and MCC by approximately 5.7%. MCC is emphasised as the primary evaluation criterion as it accounts for all four prediction outcomes simultaneously, providing a more robust measure than accuracy or F1-score when confirmed positive examples are inherently limited. Relative to Morpheus-1, true positives increased by 5.6 percentage points with only a modest rise in false positives, confirming that the gain is driven primarily by the recovery of metamorphic proteins that the secondary structure model failed to detect (Supplementary Fig. S5). Permutation importance analysis identified Mean Entropy (3Di) as the dominant predictor, with 3Di-derived features collectively outweighing secondary structure features in discriminative contribution (Fig. 2b), consistent with the existence of fold-switching proteins that maintain stable secondary structural composition while undergoing substantial rearrangement at the level of tertiary contacts. The incorporation of 3Di-derived tertiary interaction diversity within Morpheus-3D thus addresses a class of conformational plasticity that secondary structure-based approaches are inherently limited in capturing.

**Fig. 2.**
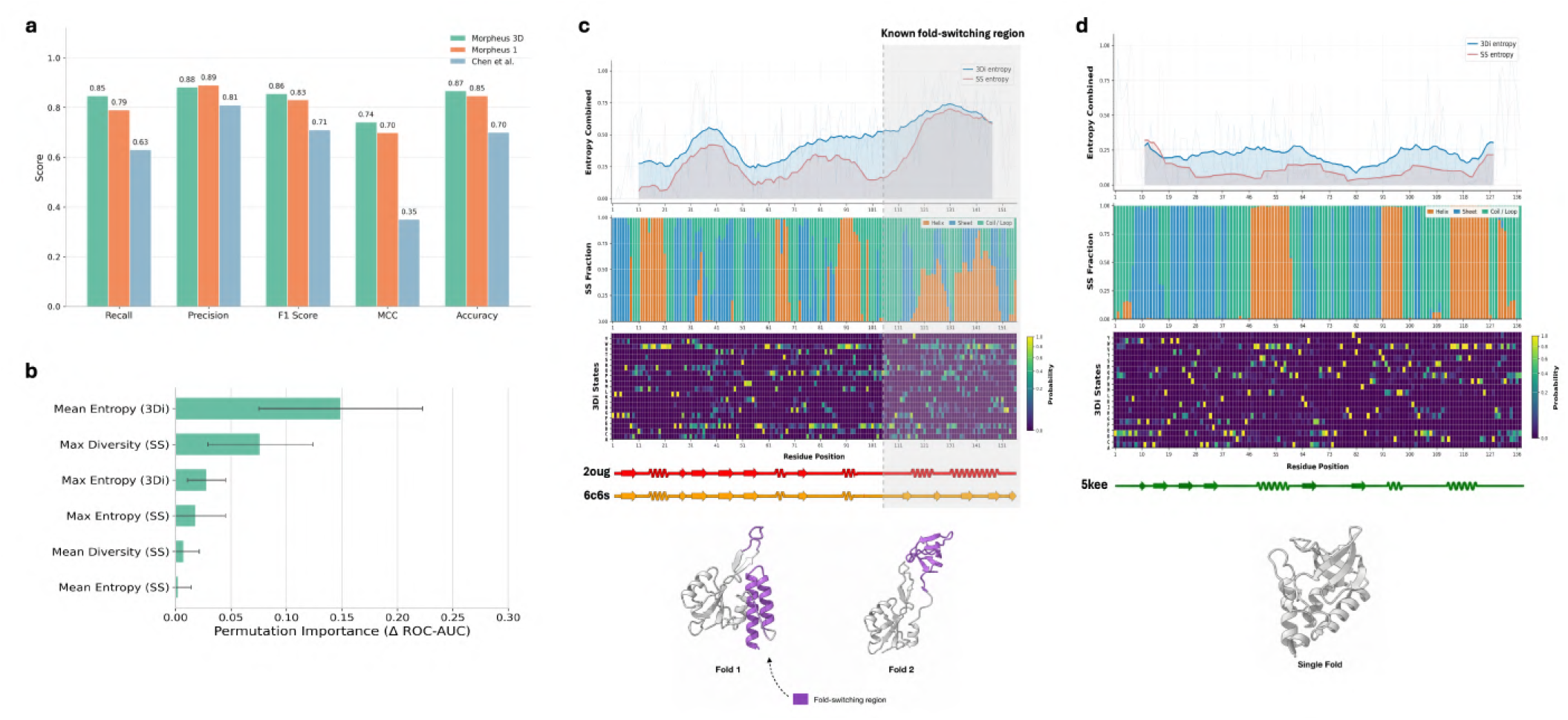
Tertiary structural diversity profiling and performance of Morpheus-3D. (**a**) Comparison of prediction performance for Morpheus-3D, Morpheus-1, and Chen et al. (**b**) Permutation feature importance showing the contribution of the six input features to model performance, reported as the mean decrease in ROC-AUC with error bars indicating standard deviation. (**c**) Residue-level structural diversity profile of the fold-switching protein RfaH (PDB: 2OUG and 6C6S). The upper panel shows rolling 3Di entropy (blue) and secondary-structure (SS) entropy (red). The middle panel shows secondary-structure propensities (orange, α-helix; blue, β-strand; green, coil/loop). The lower panel displays the probability distribution of the 20 Foldseek 3Di structural alphabet states, colored from dark purple (low probability) to yellow (high probability). Experimentally determined structures representing the two alternative folds are shown below, with the fold-switching region highlighted in purple. (**d**) Corresponding analysis for the monomorphic protein thermonuclease (PDB: 5KEE), including entropy profiles, secondary-structure propensities, 3Di state probability heatmap, and the experimentally determined structure.

### B. Residue-level entropy profiling reveals fold-switching regions within protein sequences

Beyond binary classification, a key capability of Morpheus-3D is the generation of residue-level structural diversity profiles across the full length of a query sequence, enabling spatial localisation of conformationally plastic regions rather than merely flagging a protein as metamorphic. No prior computational approach to fold-switching prediction has demonstrated this resolution. To illustrate this, entropy profiles were examined for two representative proteins: RfaH, a well-characterised metamorphic protein whose C-terminal domain undergoes a helix-to-β-barrel transition (6), and Thermonuclease, a structurally stable monomorphic protein (54). For RfaH, both *H*_3*Di*_ and *H*_*SS*_ are elevated across the sequence, with a pronounced peak in the C-terminal region spanning approximately residues 105 to 155, corresponding precisely to the domain that reorganises between an α-helical hairpin and an all-β barrel to regulate transcriptional antitermination (Fig. 2c). This is further corroborated by the 3Di diversity heatmap, which shows markedly broader and more uniform occupancy of 3Di states in this region, indicating that fragments mapping here are retrieved from structurally heterogeneous contexts. The secondary structure propensity profile for the same region displays competing helix and sheet propensities at equivalent positions, consistent with its known capacity to adopt either conformation. Thermonuclease, by contrast, displays uniformly low *H*_3*Di*_ and *H*_*SS*_ throughout, with sparse and isolated 3Di state occupancies and dominant, unambiguous secondary structure assignments at each position (Fig. 2d). To further assess the localisation capacity of Morpheus-3D, entropy profiles were examined for RfaH together with three additional experimentally characterised metamorphic proteins spanning diverse fold-switching mechanisms: Lysenin, KaiB and PimA (Fig. 3a–d). In each case, the predicted high-entropy region corresponded closely to the experimentally annotated switching region, with both *H*_3*Di*_ and *H*_*SS*_ consistently elevated across the localised fold-switching segment. In RfaH, this region coincides with the C-terminal domain whose helix-to-β-barrel transition couples RNA polymerase engagement to processive transcriptional antitermination (3, 6). Lysenin exhibits a pronounced entropy peak coinciding with the region that reorganises from a compact β-strand arrangement into an extended β-hairpin upon membrane binding, the transition that drives pore formation and membrane insertion (55). KaiB displays a well-defined entropy peak overlapping the thioredoxin-like core that switches to a ground-state fold capable of KaiC binding, the step that gates the transition from the active to the repressive phase of the circadian oscillator (7). PimA localises elevated entropy to the N-terminal domain that transitions between open and closed states upon substrate binding, a rearrangement that couples mannosyl-transferase catalysis to lipid-linked substrate accessibility, with the two states adopting distinct secondary structure configurations (56). Across all proteins, residue-level entropy mapped onto the three-dimensional structure reveals that positions with elevated *H*_3*Di*_ cluster spatially within the experimentally characterised switching regions, providing direct structural validation of the predicted localisation and demonstrating that Morpheus-3D delivers residue-level resolution directly actionable for targeted experimental characterisation.

### C. Morpheus-3D recovers new fold-switchers and identifies candidates missed by existing methods

Having established that Morpheus-3D outperforms prior classifiers and localises switching regions with residue-level precision, we next examined whether these capabilities generalise to proteins discovered independently of our training pipeline. We evaluated the model on a set of recently characterised metamorphic proteins and computationally designed two-state systems, none represented in the training data, to test whether the classifier generalises beyond its training distribution and whether its entropy profiles remain interpretable under genuinely prospective conditions. Morpheus-3D correctly classified every case tested. PopP2, a YopJ family acetyltransferase from *Ralstonia solanacearum*, undergoes an inositol hexakisphosphate (InsP6) induced α-helix to β-strand transition in its catalytic core (57). The model predicted metamorphic behaviour with high confidence and localised the switching signal to residues approximately 190– 250, corresponding precisely to the experimentally defined switching region. This localisation is reflected in the entropy trace and structural entropy maps for the two resolved conformational states (PDB: 7F3N and 5W3Y; Fig. 4a). Two additional cases were likewise correctly resolved. An ancestral double-psi β-barrel sequence experimentally shown to adopt two topologically distinct folds, a canonical DPBB and the newly described Double-Zeta β-barrel (DZBB) (58), was likewise correctly identified by Morpheus-3D, with elevated entropy overlapping the β1–β2 loop region experimentally shown to govern interconversion between the DPBB and DZBB folds (Fig. 4b). The RNA-dependent RNA polymerase nsP4 of O’nyong-nyong virus, recently characterised as a fold-switching protein (59), was similarly classified as metamorphic by the model. The most consequential prospective validation involved computationally designed two-state hinge proteins (60). These engineered systems adopt two experimentally validated conformations in response to external stimuli while largely preserving their secondary-structure composition, making them particularly challenging for secondary-structure-based approaches. Despite never being represented in the training data, Morpheus-3D correctly classified these proteins as fold-switching candidates and localized elevated entropy to the engineered hinge responsible for the conformational transition (Fig. 4c). A similar result was observed for *de novo* designed modular protein biosensors (61), where analyte binding drives transitions between closed and open conformations. Morpheus-3D correctly classified these engineered systems as fold-switching candidates despite minimal changes in secondary structure. In both cases, elevated 3Di entropy was accompanied by uniformly low secondary-structure entropy, highlighting the ability of Morpheus-3D to identify a class of conformational switching that is fundamentally inaccessible to prior methods.

### D. Proteome-wide screening uncovers extensive metamorphic potential across diverse protein families

Proteome-wide application of Morpheus-3D to 57 representative proteomes revealed that predicted metamorphic proteins are widespread across all major domains of life, although their abundance varied considerably among evolutionary lineages (Fig. 5). Functional enrichment analyses for representative species from each kingdom are provided in Supplementary Figs. S6–S12. Following disorder filtering, the fraction of predicted metamorphic proteins ranged from 1.2% to 26.5% across individual proteomes, indicating that conformational plasticity is broadly distributed but non-uniformly represented across the tree of life. Bacterial proteomes consistently showed the highest abundance of candidate metamorphic proteins, with *Mycobacterium tuberculosis* exhibiting the greatest enrichment (26.5%). Several of the species with the highest predicted content, including *M. tuberculosis, Pseudomonas aeruginosa* and *Mycobacterium leprae*, are major bacterial pathogens, suggesting an association between structural plasticity and the functional demands of a pathogenic lifestyle (62). Functional annotation further indicated that these proteins were predominantly associated with regulatory, metabolic and transport-related processes. Within Animalia, vertebrates contained substantially higher fractions of predicted metamorphic proteins than invertebrates, with *Homo sapiens* exhibiting the highest proportion (13.9%). This enrichment is consistent with the greater molecular and regulatory complexity of vertebrate proteomes, in which proteins frequently participate in extensive signalling and interaction networks; the predicted proteins were accordingly enriched in signalling, cellular communication and immune regulation. Plant proteomes, in contrast, displayed comparatively uniform prediction frequencies, with candidates enriched in pathways governing environmental sensing, stress adaptation and developmental regulation. Archaeal proteomes showed moderate levels of predicted metamorphic content. Fungal proteomes generally contained relatively few candidates, although *Saccharomyces cerevisiae* was a notable outlier, with 15.6% of its proteome predicted to possess metamorphic potential. Viral proteomes likewise displayed a high predicted frequency, whereas parasitic protists fell at intermediate levels; across these lineages, candidates were enriched primarily in regulatory and metabolic functions. Collectively, these findings indicate that predicted metamorphic proteins are pervasive across the tree of life but distributed non-randomly among evolutionary lineages. Their enrichment in bacterial pathogens, vertebrates and viruses suggests that structural plasticity is preferentially associated with organisms requiring enhanced regulatory adaptability and functional versatility.

### E. Morpheus-3D identifies evolutionary signatures associated with metamorphic proteins

To investigate the evolutionary origins of fold switching, we combined ancestral sequence reconstruction (ASR) with residue-level structural diversity profiling using Morpheus-3D. We first analysed the chemokine family, which contains one of the best-characterised evolutionary transitions to a metamorphic protein (63). Whereas human XCL1 reversibly interconverts between two distinct folded conformations (8), its close evolutionary relative CCL20 remains monomorphic (63). Consistent with previous evolutionary reconstructions, the earliest inferred ancestors of both lineages exhibited uniformly low 3Di and secondary-structure entropy, indicative of structurally constrained proteins. Along the XCL1 lineage, however, a marked increase in entropy was observed at the Anc.2 to Anc.3 transition, followed by a progressive increase towards modern XCL1 (Fig. 6a,b). In contrast, entropy remained consistently low throughout the CCL20 lineage (Fig. 6c). These observations closely mirror the experimentally proposed evolutionary acquisition of fold switching in XCL1, while indicating that the CCL20 lineage retained a single stable fold (63). To determine whether this pattern extends beyond chemokines, we next examined KaiB, a circadian clock protein that undergoes reversible fold switching (7). As observed for XCL1, reconstructed ancestral KaiB proteins displayed uniformly low entropy profiles, whereas modern proteins exhibited pronounced localised increases in 3Di entropy (Fig. 6d,e). Notably, the entropy maximum coincided with the experimentally characterised switching region (7). Mapping amino acid substitutions onto the phylogeny further showed that mutations accumulated predominantly within this same segment, while the remainder of the protein remained comparatively conserved, linking the emergence of conformational plasticity to sequence diversification within the switching region (Fig. 6f). Finally, we investigated the evolutionary architecture of the RfaH/NusG superfamily. Unlike XCL1, which followed a single evolutionary transition from a monomorphic ancestor (63), predicted metamorphic proteins were distributed across multiple independent branches of the RfaH/NusG phylogeny (Supplementary Fig. S13). This pattern is consistent with previous studies proposing a polyphyletic origin of fold switching within the super-family (64), suggesting repeated and independent acquisition of conformational plasticity during evolution. Together, these analyses demonstrate that Morpheus-3D captures evolutionary signatures associated with the emergence of fold switching across diverse protein families. Beyond identifying metamorphic proteins, the framework reveals lineage-specific changes in structural diversity, providing a means to trace the evolutionary acquisition of conformational plasticity through reconstructed ancestral sequences. This capacity to trace lineage-specific change also suggests a hypothesis about the ordering of variation across levels. Since evolution acts directly on sequence, any change at the structural level must originate there: mutations may first perturb local tertiary packing, captured as a rise in 3Di entropy, before this is expressed as a change in secondary-structure entropy, since the 3Di alphabet encodes a finer-grained structural description than the three-state SS classification and may be more sensitive to early, small-scale perturbation. We do not test this ordering directly here, and confirming it would require examining whether 3Di entropy consistently precedes SS entropy across ancestral nodes; we note it as a testable hypothesis rather than a conclusion of the present analysis.

### F. Shortlisted Candidate Proteins for Experimental Follow-Up

Although proteome-wide application of Morpheus-3D identified numerous proteins with predicted metamorphic potential, experimental validation remains inherently low throughput and therefore requires careful candidate prioritisation. To identify the most promising targets, we applied a stringent filtering framework requiring: (i) a Morpheus-3D confidence score greater than 0.60; (ii) overlap between the predicted fold-switching (FS) region and a UniProt domain annotation (40); (iii) less than 50% predicted disorder within the putative switching region; and (iv) the presence of both α-helical and β-sheet propensities within the predicted switching segment, consistent with the potential to adopt alternative secondary structures. Application of these filters yielded ten high-confidence candidates spanning diverse taxonomic groups and functional classes (Table 1; Supplementary Figs. S14–S18). Several predicted switching regions overlap domains previously implicated in structural plasticity or conformational rearrangements, providing independent support for the biological relevance of the predictions. Among the highest-confidence candidates, NusG from *Salmonella typhimurium* (UniProt: P0AA02) contains a predicted switching region within a protein family recently shown to include multiple sequence-diverse members capable of reversible α-helix-to-β-sheet transitions (65). This observation is consistent with our evolutionary analysis of the RfaH/NusG superfamily, in which predicted metamorphic proteins were distributed across multiple independent evolutionary lineages. Likewise, the transcriptional regulator CmeR from *Campylobacter jejuni* (UniProt: Q0PBE2) contains a predicted high-entropy region within its N-terminal DNA-binding domain that overlaps the unusually flexible segment replacing the canonical TetR recognition helix, a region proposed to undergo a coil to helix transition during DNA binding (66). In human HSPB2 (UniProt: Q16082), the predicted fold-switching region lies within the conserved α-crystallin domain, which mediates dimerization and oligomeric assembly in small heat shock proteins (67). Collectively, the concordance between Morpheus-3D predictions and previously characterised structurally dynamic regions highlights these proteins as compelling candidates for future biophysical characterisation and for expanding the currently known repertoire of metamorphic proteins.

**Table 1.** Shortlisted high-confidence candidate metamorphic proteins for experimental validation.

| UniProt ID | Protein name | Species | Confidence score | Length (aa) | Predicted FS region |
| --- | --- | --- | --- | --- | --- |
| P0AA02 | Transcription termination/antitermination protein NusG | *Salmonella typhimurium* | 0.87 | 181 | 40–66 |
| A4HVI6 | 40S ribosomal protein S12 | *Leishmania infantum* | 0.87 | 141 | 82–112 |
| Q0PBE2 | Transcriptional regulator CmeR | *Campylobacter jejuni* | 0.65 | 210 | 38–57 |
| P06110 | Chemotaxis protein CheW | *Salmonella typhimurium* | 0.88 | 167 | 90–142 |
| Q16082 | Heat shock protein beta-2 (HSPB2) | *Homo sapiens* | 0.67 | 182 | 86–101 |
| Q9UHD4 | Lipid transferase CIDEB | *Homo sapiens* | 0.65 | 219 | 49–79 |
| K0F3Y5 | Deoxyuridine 5'-triphosphate nucleotidohydrolase | *Nocardia brasiliensis* | 0.66 | 163 | 42–52 |
| K7W064 | Glutaredoxin domain-containing protein | *Zea mays* | 0.87 | 159 | 105–131 |
| I6X9X6 | Methyltransferase domain-containing protein | *Mycobacterium tuberculosis* | 0.66 | 270 | 45–69 |
| Q4E4Q8 | Large ribosomal subunit protein uL15 | *Trypanosoma cruzi* | 0.84 | 145 | 77–101 |

### G. The Morpheus-3D web platform enables interactive exploration of predictions

To make residue-level predictions directly explorable without requiring local installation, we implemented Morpheus-3D as an interactive web platform (https://morpheus.slicearrow.com/) (Fig. 7; see Methods). Because the underlying *H*_3*Di*_ and *H*_*SS*_ profiles are computed directly from sequence, a single prediction can be mapped onto a structure whose sequence matches the query via alignment; for a metamorphic protein, this includes each of its alternative experimentally resolved conformations, since these share the same or near-identical sequence. We illustrate this for Lysenin, a pore-forming toxin whose fold-switching mechanism is described above (Fig. 3b): supplying the PDB accession of the compact pre-pore conformation independently computes and maps the entropy profile onto that structure (Fig. 7b), and supplying the accession of the extended membrane-inserted conformation independently computes and maps the corresponding profile onto the alternative fold (Fig. 7c); because both queries derive from the same underlying sequence, elevated entropy consistently localizes to the same switching region in both cases. The full retrieval and rendering workflow, including the sequence-to-model matching logic, is summarized in Supplementary Fig. S19.

## Scope and Limitations

Morpheus-3D is designed as a sequence-based screening and localization tool, and its predictions should be interpreted within that scope. The model identifies candidate fold-switching proteins and their putative switching regions; it does not predict the alternative structural states themselves, nor does it provide mechanistic or kinetic information about how or under what conditions a transition occurs. A central limitation stems from training data scarcity: fewer than a hundred experimentally validated metamorphic proteins are currently known, constraining both the diversity of switching mechanisms captured during training and our ability to rigorously estimate generalisation error. The 3Di-based entropy framework is also sensitive to the composition of the underlying structural database; fragments from sparsely represented protein families yield less reliable diversity estimates than those from well-populated structural neighbourhoods, which may bias detection toward families with greater structural coverage in the Protein Data Bank and AlphaFold2 predictions. Disorder filtering using IUPred and MetaPredict mitigates but does not eliminate false positives arising from intrinsically disordered regions, which can superficially resemble conformational plasticity in entropy profiles. Additionally, because the approach relies on fragment-level retrieval against a reference database, its sensitivity may be reduced for proteins with limited structural homology to characterised folds, including some lineage-specific or rapidly evolving proteins. The proteome-wide enrichment patterns reported here are correlative and descriptive; while they motivate hypotheses linking structural plasticity to pathogenicity, regulatory complexity, and environmental adaptation, they do not establish causal relationships. Finally, the shortlisted candidates emerging from our filtering framework still require experimental structural characterisation, as computational prediction alone cannot confirm the existence or functional relevance of an alternative fold. We view these limitations not as barriers but as a roadmap for future work, particularly the expansion of experimentally validated training data and tighter integration with structure-based and biophysical validation pipelines.

**Fig. 3.**
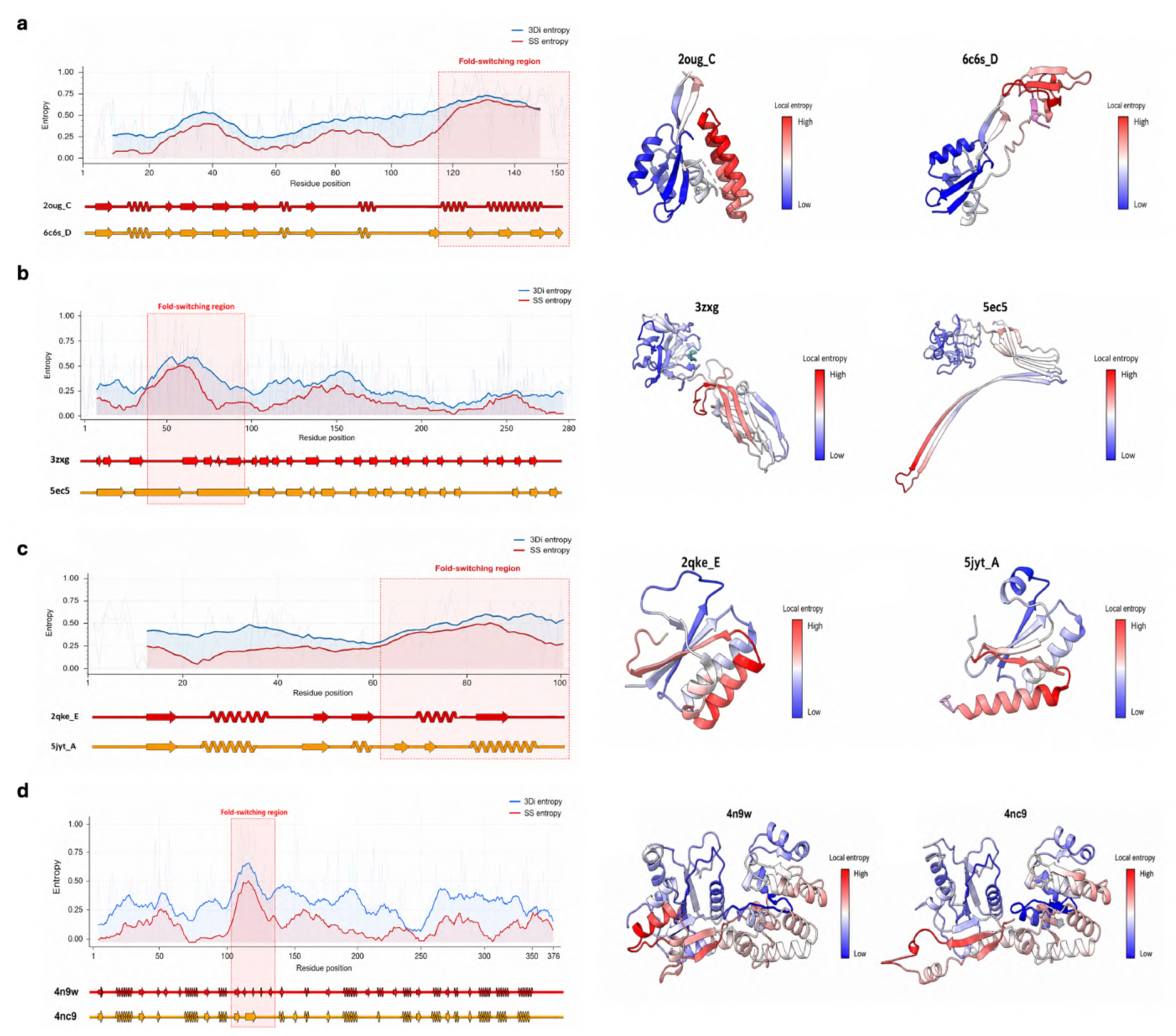
Residue-level localization of experimentally validated fold-switching regions in diverse metamorphic proteins. (**a–d**) Residue-level structural diversity profiles (left) and corresponding mapping of 3Di entropy onto experimentally determined structures (right) for four fold-switching proteins. The left panels show rolling 3Di entropy (blue) and secondary-structure (SS) entropy (red). Experimentally characterized fold-switching regions are highlighted by the red shaded boxes. Secondary-structure annotations derived from the corresponding protein structures are shown below each profile. The right panels display 3Di entropy mapped onto the protein structures, colored from blue (low 3Di entropy) to red (high 3Di entropy). (**a**) RfaH (PDB: 2OUG and 6C6S). (**b**) Lysenin (PDB: 3ZXG and 5EC5). (**c**) KaiB (PDB: 2QKE and 5JYT). (**d**) PimA (PDB: 4N9W and 4NC9).

**Fig. 4.**
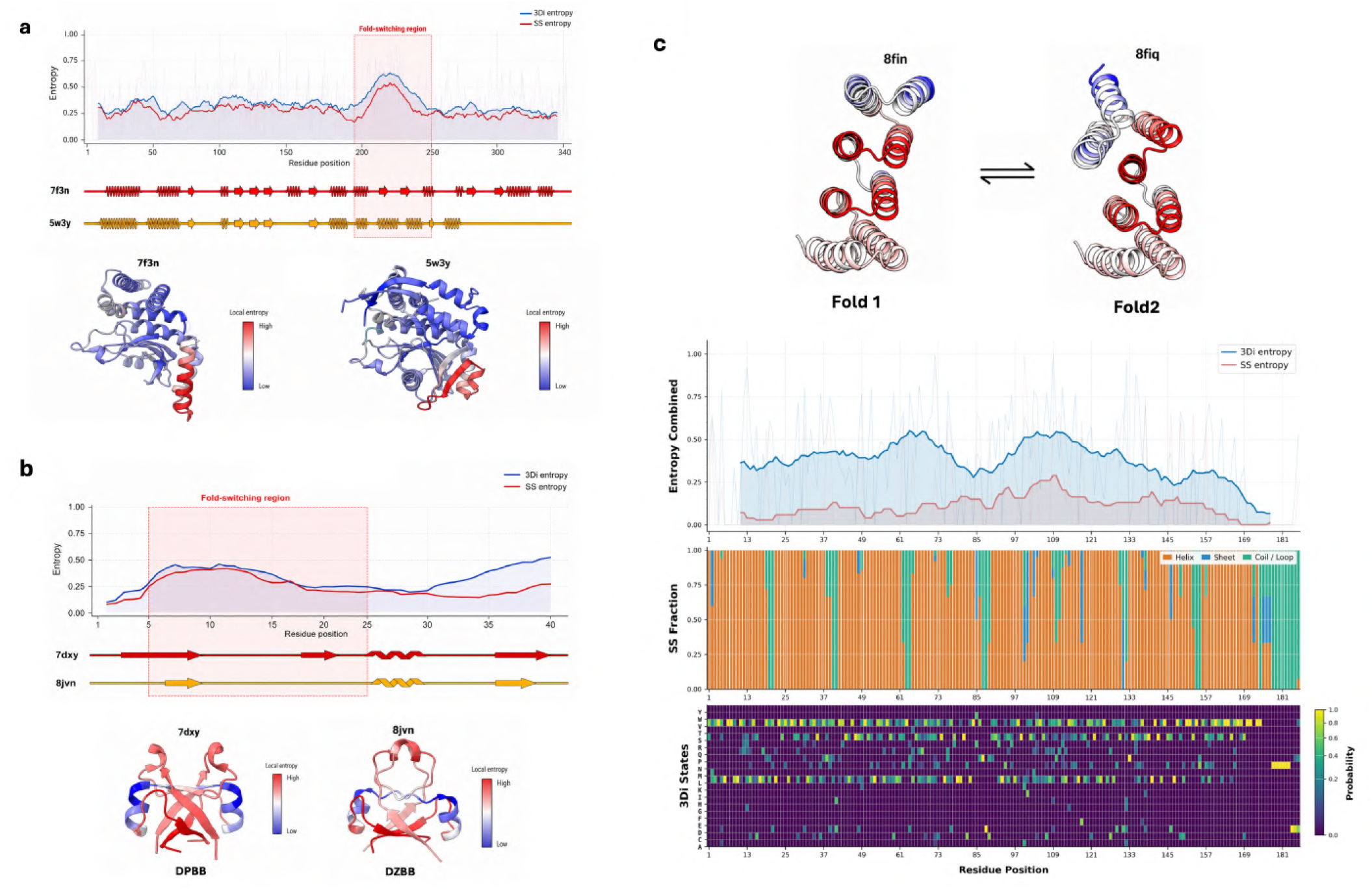
Validation of Morpheus-3D on recently characterized and engineered fold-switching proteins. (**a**) PopP2 showing the residue-level 3Di entropy (blue) and secondary-structure (SS) entropy (red) profiles, corresponding secondary-structure annotations, and 3Di entropy mapped onto the two experimentally determined conformations (PDB: 7F3N and 5W3Y), colored from blue (low) to red (high). The experimentally characterized fold-switching region is indicated by the red shaded box. (**b**) Equivalent analysis for the ancestral DPBB/DZBB fold-switching protein (PDB: 7DXY and 8JVN). (**c**) Computationally designed two-state hinge protein showing the experimentally determined Fold 1 and Fold 2 conformations (PDB: 8FIN and 8FIQ), together with the corresponding 3Di entropy (blue) and SS entropy (red) profiles, secondary-structure propensity plot (orange, α-helix; blue, β-strand; green, coil/loop), and the probability distribution of the 20 Foldseek 3Di structural alphabet states, shown from dark purple (low probability) to yellow (high probability).

**Fig. 5.**
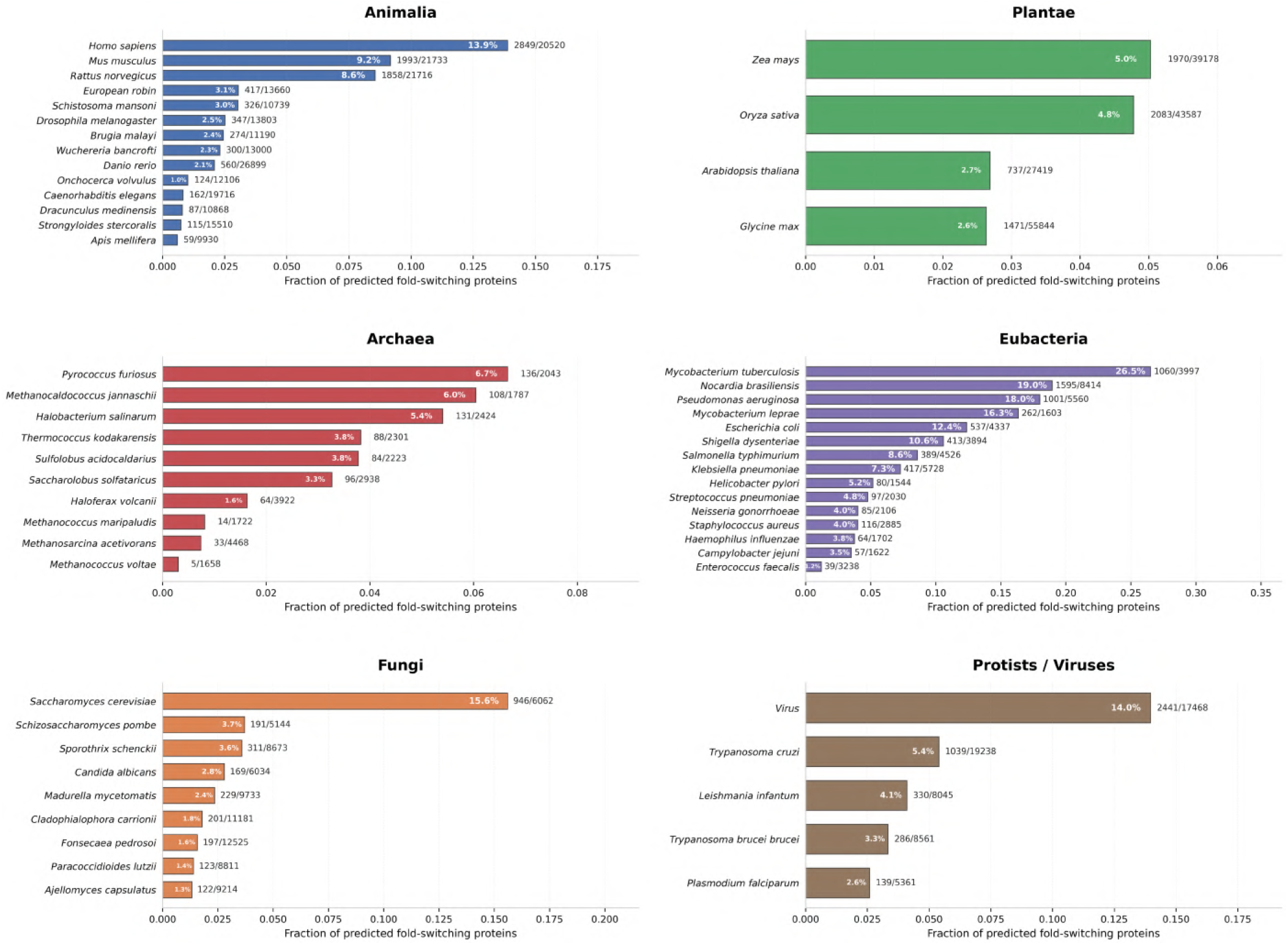
Proteome-wide distribution of predicted fold-switching proteins across representative proteomes. Horizontal bar plots showing the fraction of proteins predicted to exhibit fold-switching behavior across 57 representative proteomes spanning six major taxonomic groups: Animalia (blue), Plantae (green), Archaea (red), Eubacteria (purple), Fungi (orange), and Protists/Viruses (brown). Species within each kingdom are ranked by the fraction of predicted fold-switching proteins. Percentages indicate the proportion of predicted fold-switching proteins, and values adjacent to each bar denote the number of predicted fold-switching proteins relative to the total proteome size.

**Fig. 6.**
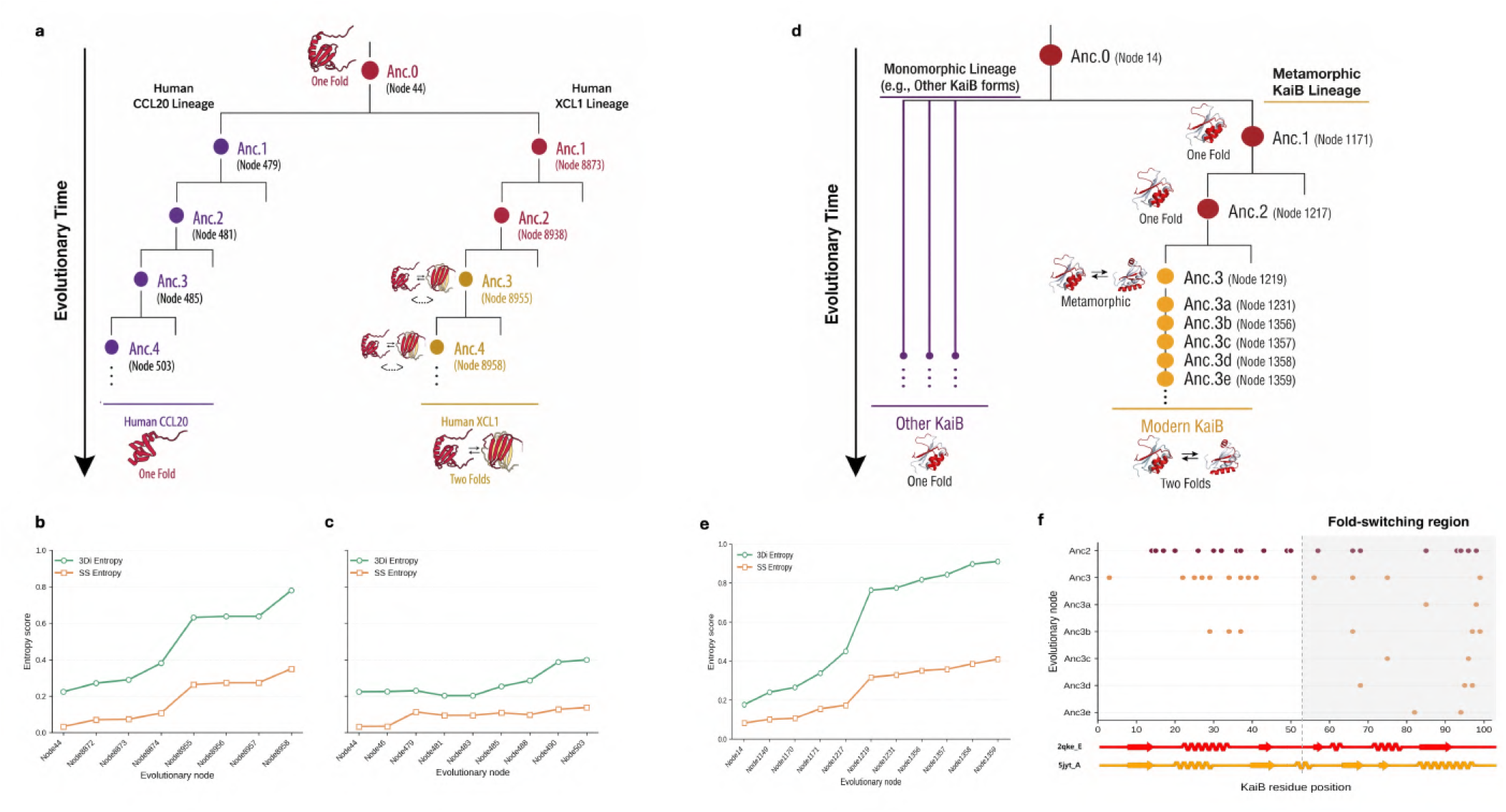
Evolutionary emergence of fold switching revealed by ancestral sequence reconstruction. (**a**) Reconstructed evolutionary trajectories of the XCL1 and CCL20 lineages illustrating the inferred emergence of fold switching. (**b**,**c**) Residue-level 3Di entropy (green) and secondary-structure (SS) entropy (orange) across reconstructed ancestral nodes for the XCL1 and CCL20 lineages, respectively. (**d**) Reconstructed evolutionary trajectory of the KaiB lineage showing the transition from monomorphic to metamorphic proteins. (**e**) Residue-level 3Di (green) and SS (orange) entropy across reconstructed KaiB ancestors. (**f**) Distribution of amino acid substitutions across KaiB ancestral nodes, mapped onto the experimentally characterized fold-switching region, with secondary structures from the two conformational states (PDB: 2QKE and 5JYT) shown below.

**Fig. 7.**
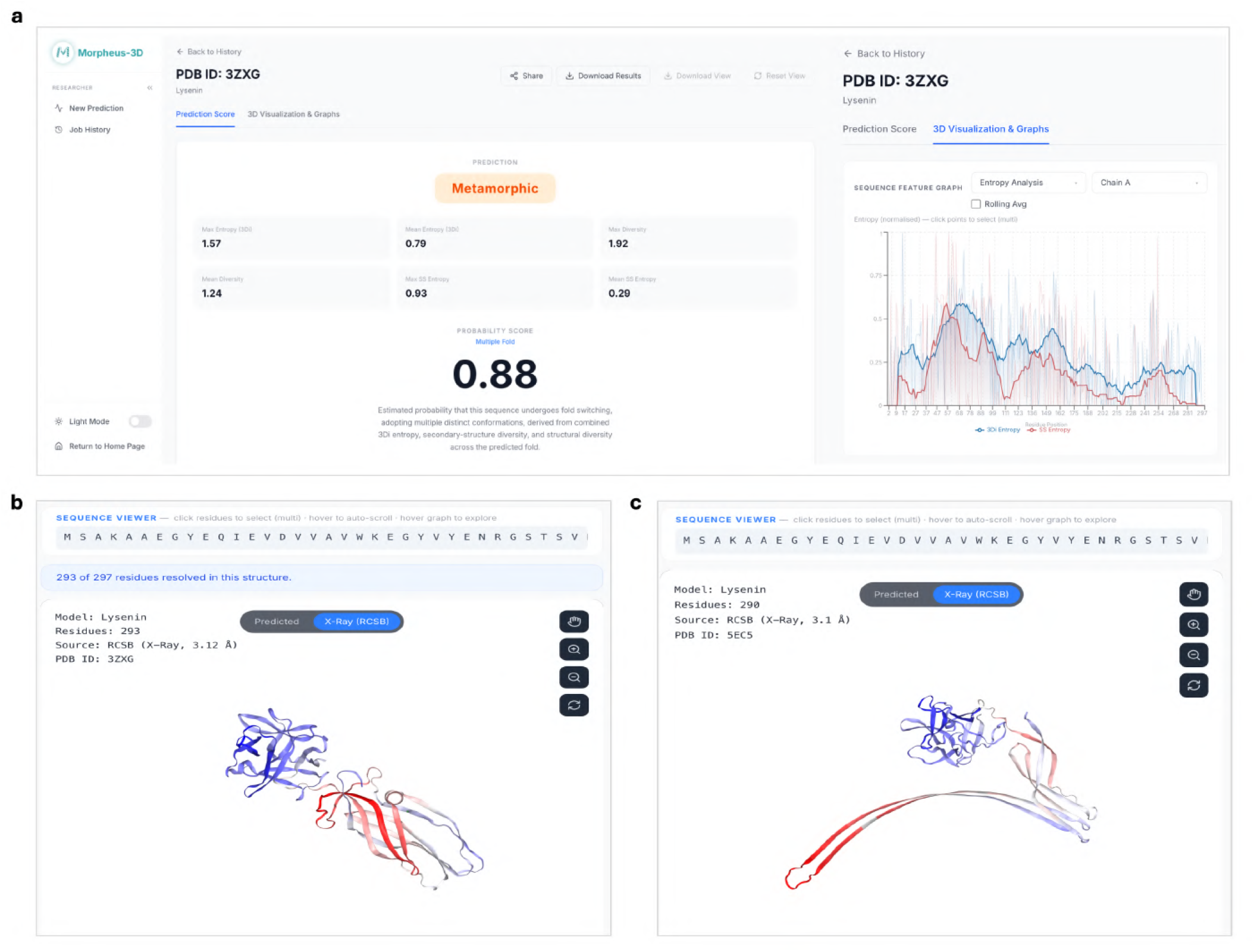
The Morpheus-3D interactive web platform, illustrated for Lysenin. (**a**) Prediction summary and sequence feature graph. The left panel shows the classification type (Metamorphic), summary diversity metrics, and probability score (0.88), computed once from the input sequence. The right panel shows rolling 3Di entropy (blue) and SS entropy (red) against residue positions; the N-terminal peak matches the switching segment in Fig. 3b. (**b, c**) The same entropy profile mapped, via PDB accession, onto the two alternative conformations of Lysenin: compact pre-pore (PDB: 3ZXG) (**b**) and extended, membrane-inserted (PDB: 5EC5) (**c**), coloured blue (low) to red (high) 3Di entropy. Elevated entropy localizes to the same *β*-hairpin in both folds, showing that a single sequence-based prediction identifies the switching region independent of the mapped structure. All panels are interactive; results are exportable from the interface.

## Conclusion

We present Morpheus-3D, a sequence-based framework for identifying fold-switching proteins while simultaneously localizing the specific regions responsible for conformational plasticity. By quantifying residue-level structural diversity through entropy profiles derived from a structural alphabet representation, Morpheus-3D extends beyond conventional binary classification to provide spatially resolved predictions of fold-switching regions directly from sequence. This residue-level capability distinguishes Morpheus-3D from previous computational approaches and enables experimentally actionable predictions. Benchmarking against experimentally characterized metamorphic proteins demonstrated that Morpheus-3D accurately recovers known switching regions, including the C-terminal helix-to-β-barrel transition of RfaH, while producing uniformly low-entropy profiles for structurally stable monomorphic proteins. Importantly, the framework generalized to proteins that were discovered independently of its training data, correctly identifying recently characterized natural metamorphic proteins such as PopP2 and the O’nyong-nyong virus RNA-dependent RNA polymerase nsP4, as well as computationally designed two-state proteins, including stimulus-responsive hinge proteins and modular biosensors. These engineered systems undergo substantial tertiary structural rearrangements with minimal changes in secondary structure, making them difficult to detect using secondary-structure-based approaches. The successful identification of these proteins demonstrates that Morpheus-3D captures structural plasticity that is fundamentally inaccessible to earlier methods. Application of Morpheus-3D to 57 representative proteomes spanning six biological kingdoms further revealed that predicted metamorphic proteins are widespread across the tree of life. Elevated frequencies were observed in evolutionarily adaptable and pathogenic organisms, particularly *Mycobacterium tuberculosis*, while proteins involved in signalling and regulatory processes were enriched within Animalia. These observations suggest that conformational plasticity represents an evolutionarily conserved strategy for expanding functional versatility in organisms facing complex regulatory or environmental challenges. Evolutionary analyses integrating ancestral sequence reconstruction with entropy profiling further showed that Morpheus-3D captures the emergence of conformational plasticity over evolutionary time. Distinct evolutionary trajectories were observed for XCL1 and CCL20, while analyses of the RfaH/NusG superfamily revealed a polyphyletic distribution of fold-switching behaviour, indicating that metamorphism has arisen multiple times independently during evolution. Collectively, these results establish Morpheus-3D as a scalable framework for discovering metamorphic proteins directly from sequence, without requiring experimentally determined structures. By combining accurate prediction with residue-level localization, the method substantially narrows the experimental search space for validating fold-switching regions and prioritizes high-confidence candidates across diverse organisms and functional classes. To make these residue-level predictions directly usable, Morpheus-3D is accompanied by an interactive web platform that maps the entropy profiles and predicted fold-switching regions onto the three-dimensional structure. By linking the sequence, the profiles and the structure in a single view, the platform allows predicted switching regions to be examined in their structural context and prioritized for experimental characterization. Beyond facilitating the discovery of previously unrecognized metamorphic proteins, Morpheus-3D provides a platform for investigating the evolutionary origins of structural plasticity and offers new opportunities for protein engineering, rational protein design, and therapeutic discovery.

## Data Availability

The Morpheus-3D source code, trained models, and documentation are openly available at GitHub (https://github.com/codesrivastavalab/Morpheus-3D). The Morpheus-3D database, comprising the sequence archive and indexed SQLite database used for fragment retrieval and structural diversity profiling, is publicly available through Hugging Face (https://huggingface.co/datasets/sreeharshk/Morpheus3D-Database). These repositories contain all resources required to reproduce the analyses reported in this study.

## ACKNOWLEDGEMENTS

AS acknowledges the financial support from the Indian Institute of Science (IISc) and the high-performance computing facility “Beagle” that was set up from grants by the erstwhile IISc-DBT partnership programme. SK and AS also acknowledge the computing facilty from SERC, IISc. AS also acknowledges the FIST program sponsored by the Department of Science and Technology, India that supports the MBU infrastructure. AS would also like to thank the Teams Science Grant from the DBT-Wellcome Trust India Alliance (Grant number: IA/TSG/21/1/600245). AS also thanks the DBT National Network Project (NNP) grant (BT/PR40323/BTIS/137/78/2023), the Matrics grants (MTR/2023/001040) and the ARG grant (ANRF/ARG/2025/009141/LS)from ANRF, India. SK and AS thank Prof. Karthish Manthiram from Caltech for discussions during his sabbatical stay at MBU as a part of his Infosys Prize fellowship visit to India.

AA and AL thank Shakeera Begum, who led the front-end development, and Anusha Bobili and Raju C. for their assistance with the front-end implementation. AA and AL are grateful to Sajid Hussain for leading the back-end development and to Jobin A. J. and Srinidhi Srikanth for DevOps support. We also acknowledge Sirpi Products and Services Private Limited for hosting the Morpheus-3D website on their in-house platform, Slicearrow (68).

## Supplementary Information

### Supplementary Note 1: Figures

**Fig. S1.**
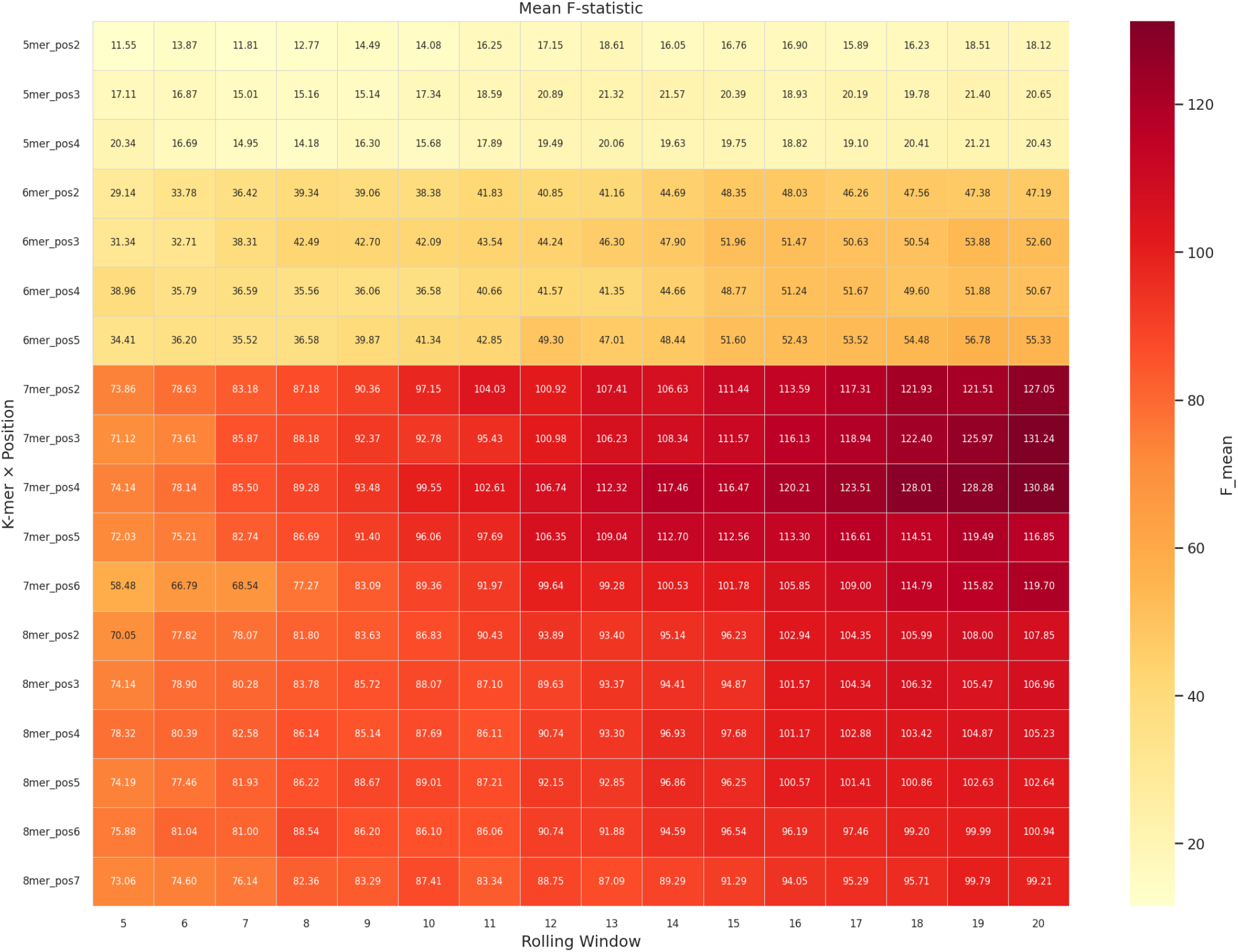
Balanced one-way ANOVA for optimization of Morpheus-3D parameters. Mean F-statistics obtained from balanced one-way ANOVA using the maximum rolling 3Di entropy are shown for different fragment lengths, representative residue positions, and rolling-window sizes. Warmer colours indicate stronger discrimination between metamorphic and monomorphic proteins. The highest F-statistics were observed for 7-mer fragments, representative positions 3–4, and larger rolling windows, supporting the selection of a 7-mer fragment, position 3, and a 20-residue rolling window for subsequent analyses.

**Fig. S2.**
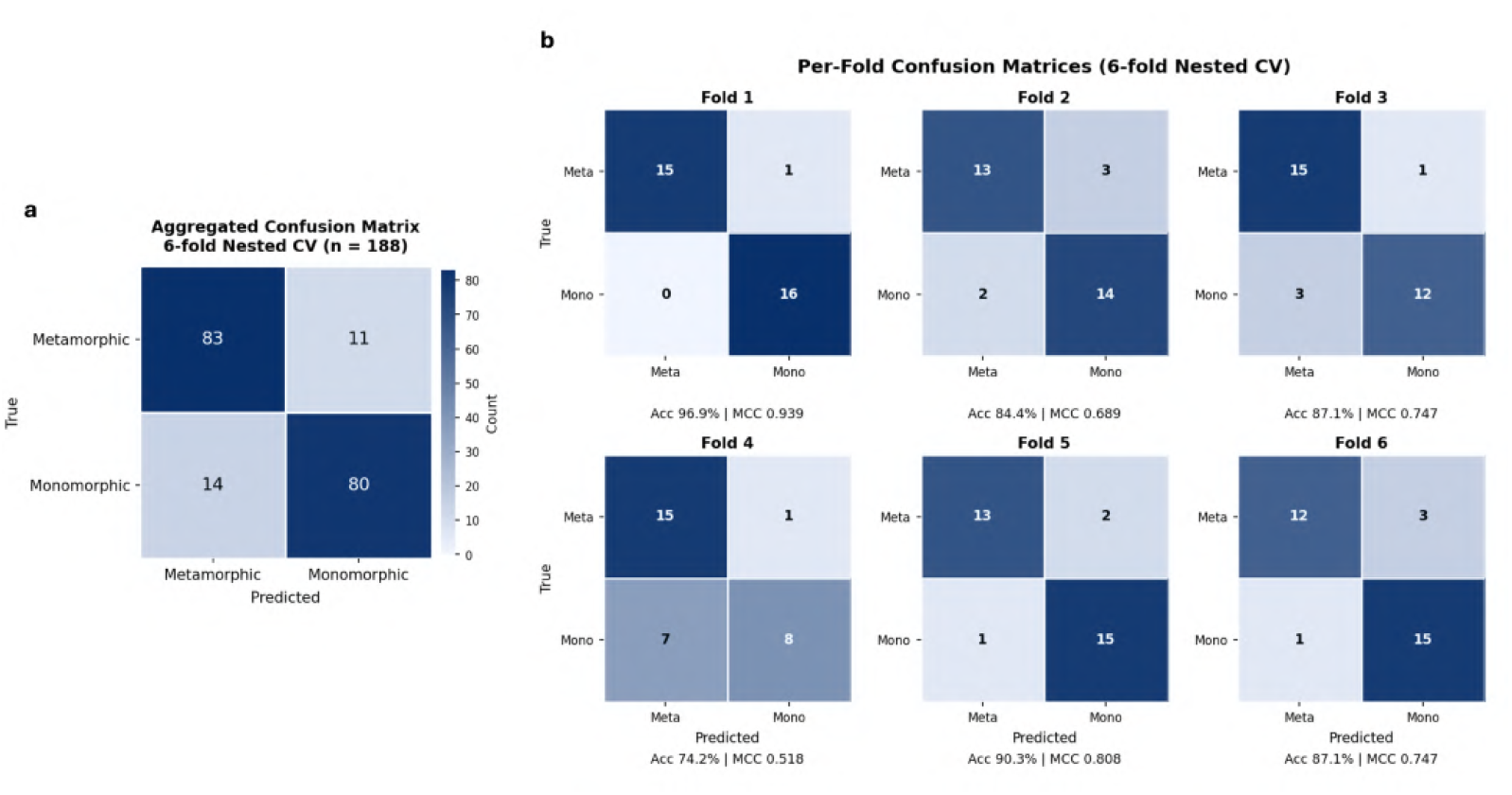
Confusion matrices of the final Morpheus-3D classifier. (**a**) Aggregated confusion matrix obtained across the six outer folds of the nested cross-validation. (**b**) Confusion matrices for each individual outer fold, with the corresponding accuracy and Matthews correlation coefficient (MCC). The results demonstrate consistent classification performance across cross-validation folds.

**Fig. S3.**
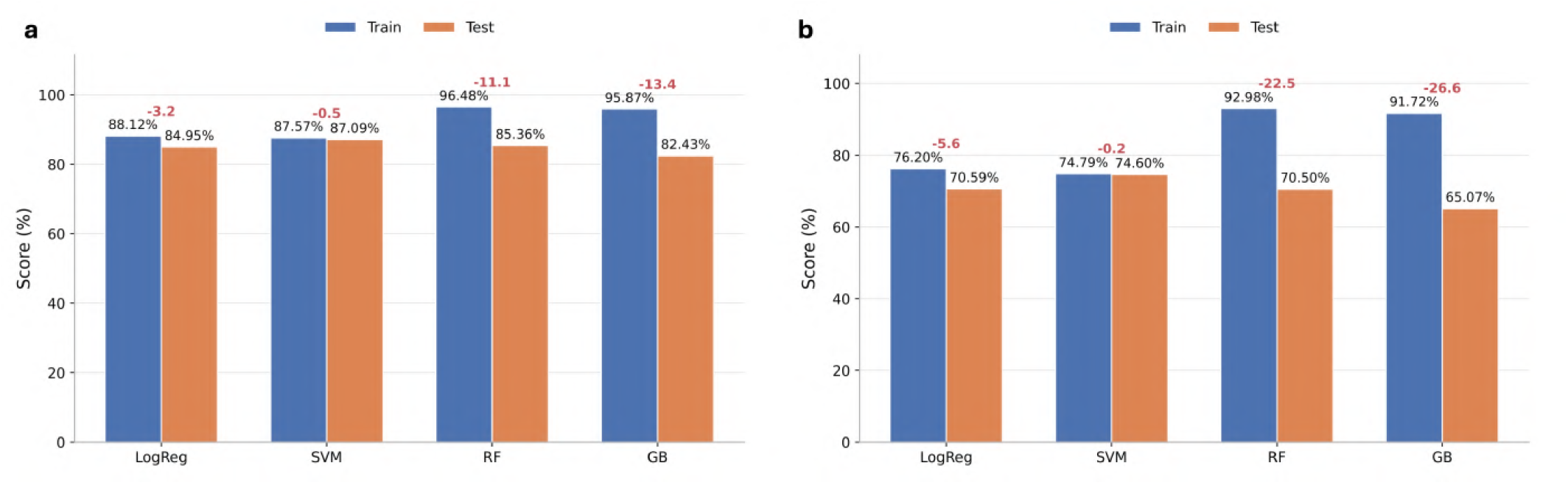
Training and test performance of candidate machine-learning models. (**a**) Training and test accuracies of Logistic Regression (LogReg), Support Vector Machine (SVM), Random Forest (RF), and Gradient Boosting (GB). (**b**) Corresponding Matthews correlation coefficients (MCC). SVM exhibited the smallest train–test performance gap while maintaining competitive predictive performance, supporting its selection as the final classifier for Morpheus-3D.

**Fig. S4.**
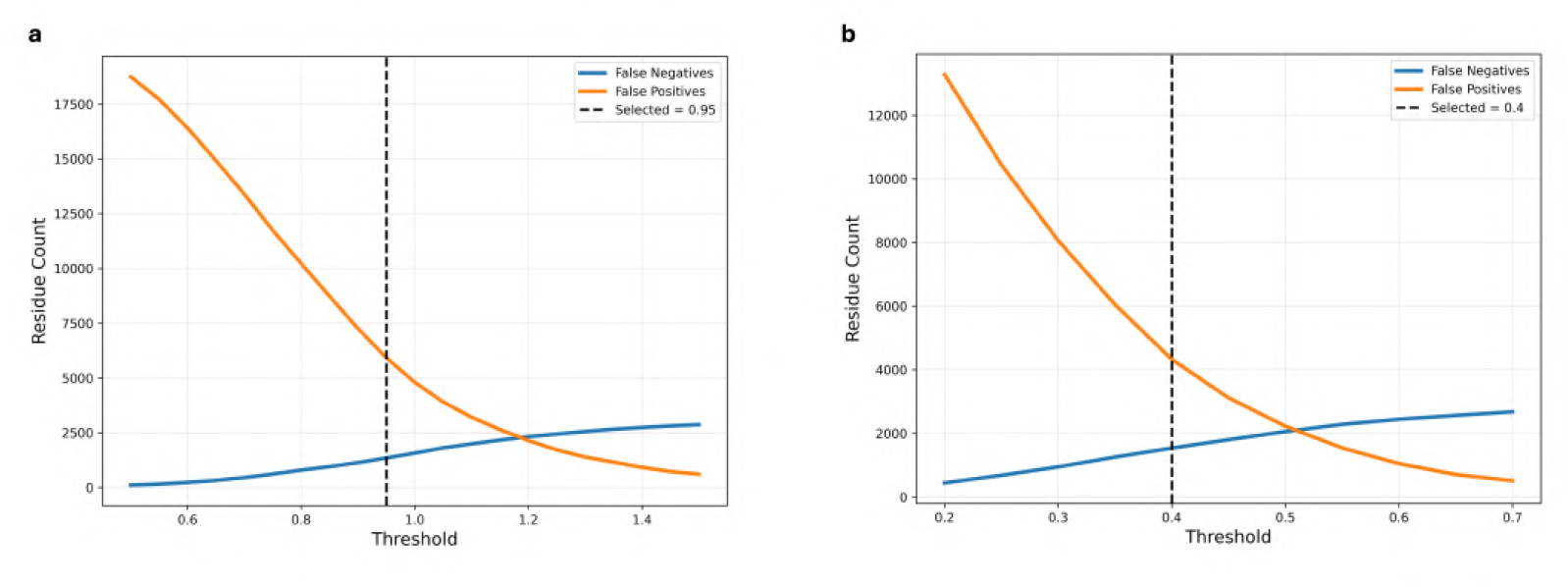
Selection of residue-level entropy thresholds for fold-switching region identification. (**a**) False-positive (orange) and false-negative (blue) residue counts obtained across a range of 3Di entropy thresholds using the benchmark dataset of experimentally annotated fold-switching proteins. (**b**) Equivalent analysis for secondary-structure entropy thresholds. Dashed lines indicate the selected thresholds (3Di entropy = 0.95; secondary-structure entropy = 0.40), which were subsequently used for residue-level identification of putative fold-switching regions.

**Fig. S5.**
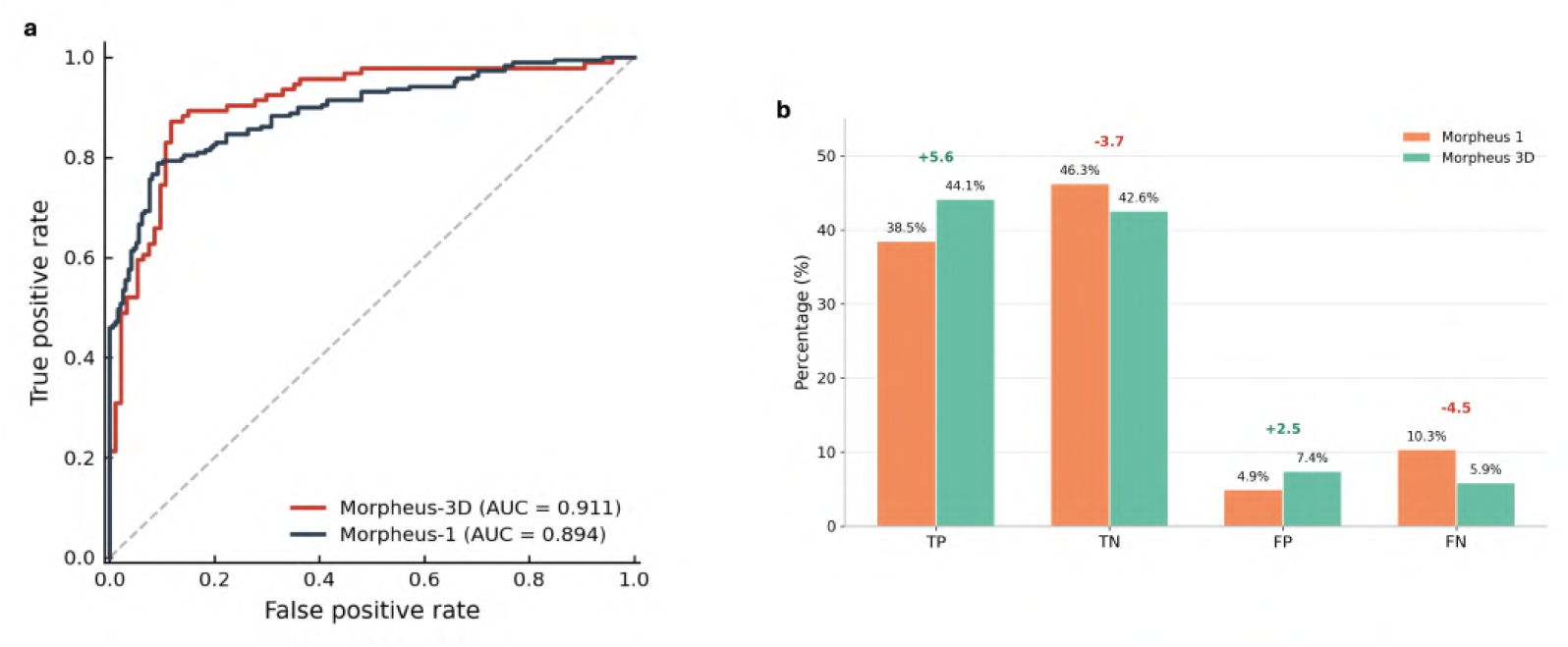
Performance comparison between Morpheus-3D and Morpheus-1. (**a**) Receiver operating characteristic (ROC) curves comparing the predictive performance of Morpheus-3D and Morpheus-1. (**b**) Distribution of true positives (TP), true negatives (TN), false positives (FP), and false negatives (FN) across the benchmark dataset. Morpheus-3D improves the recovery of metamorphic proteins while reducing false-negative predictions relative to Morpheus-1.

**Fig. S6.**
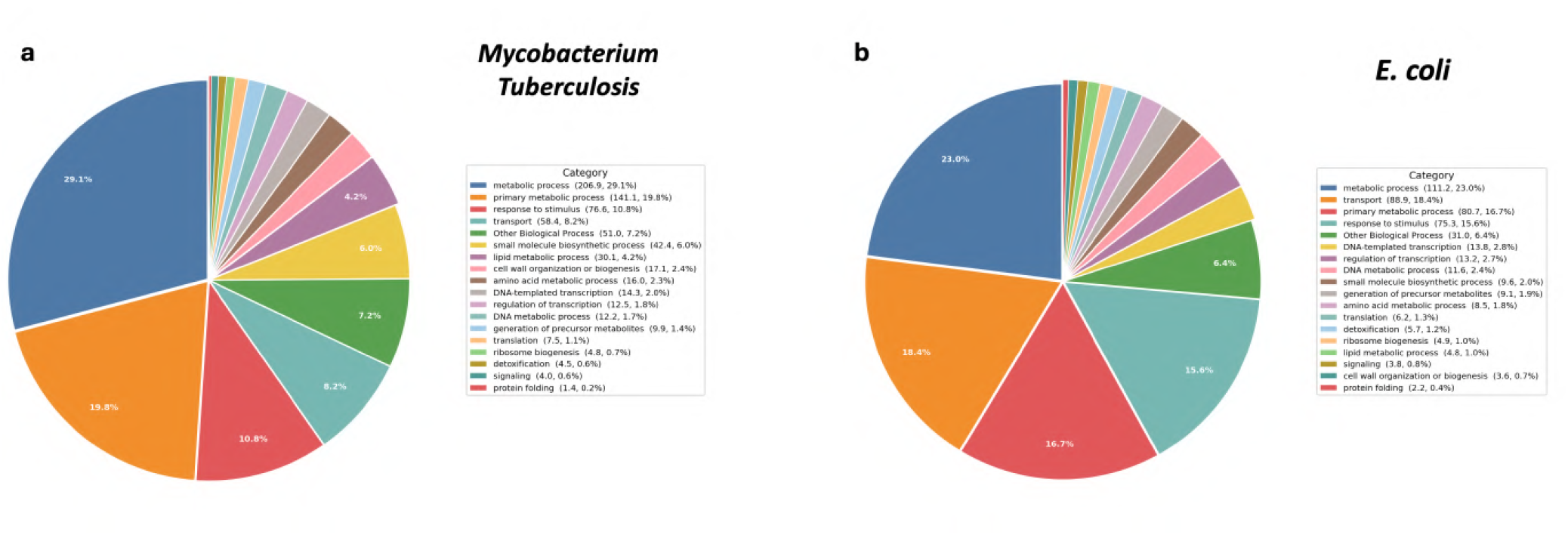
Gene Ontology analysis of predicted metamorphic proteins in representative eubacterial proteomes. Distribution of high-level Gene Ontology (GO) biological process categories among predicted metamorphic proteins in (**a**) *Mycobacterium tuberculosis* and (**b**) *Escherichia coli*. GO terms were grouped into high-level biological process categories to facilitate comparison of the functional landscape of predicted metamorphic proteins across representative eubacterial species.

**Fig. S7.**
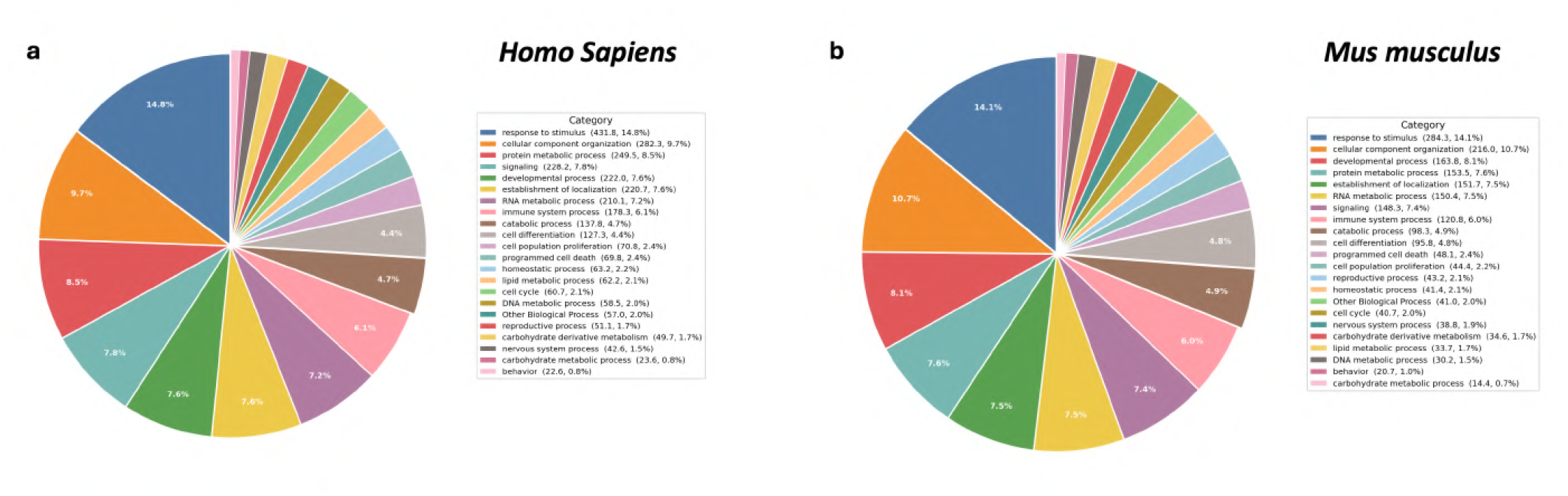
Gene Ontology analysis of predicted metamorphic proteins in representative animal proteomes. Distribution of high-level Gene Ontology (GO) biological process categories among predicted metamorphic proteins in (**a**) *Homo sapiens* and (**b**) *Mus musculus*. GO terms were grouped into high-level biological process categories to facilitate comparison of the functional landscape of predicted metamorphic proteins across representative animal species.

**Fig. S8.**
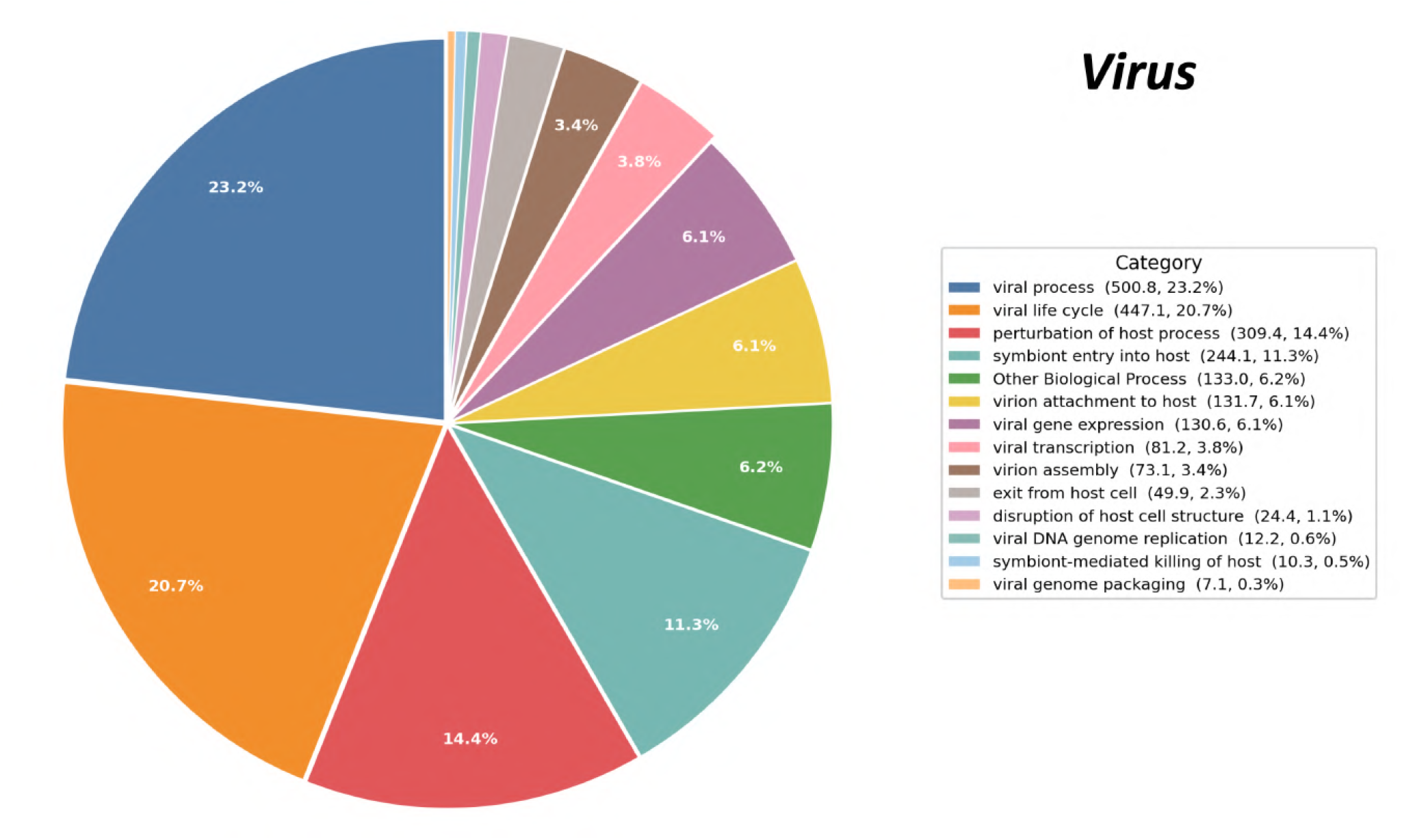
Gene Ontology analysis of predicted metamorphic proteins in the viral proteome. Distribution of high-level Gene Ontology (GO) biological process categories among predicted metamorphic viral proteins. GO terms were grouped into high-level biological process categories to summarize the functional landscape of predicted metamorphic proteins in viruses.

**Fig. S9.**
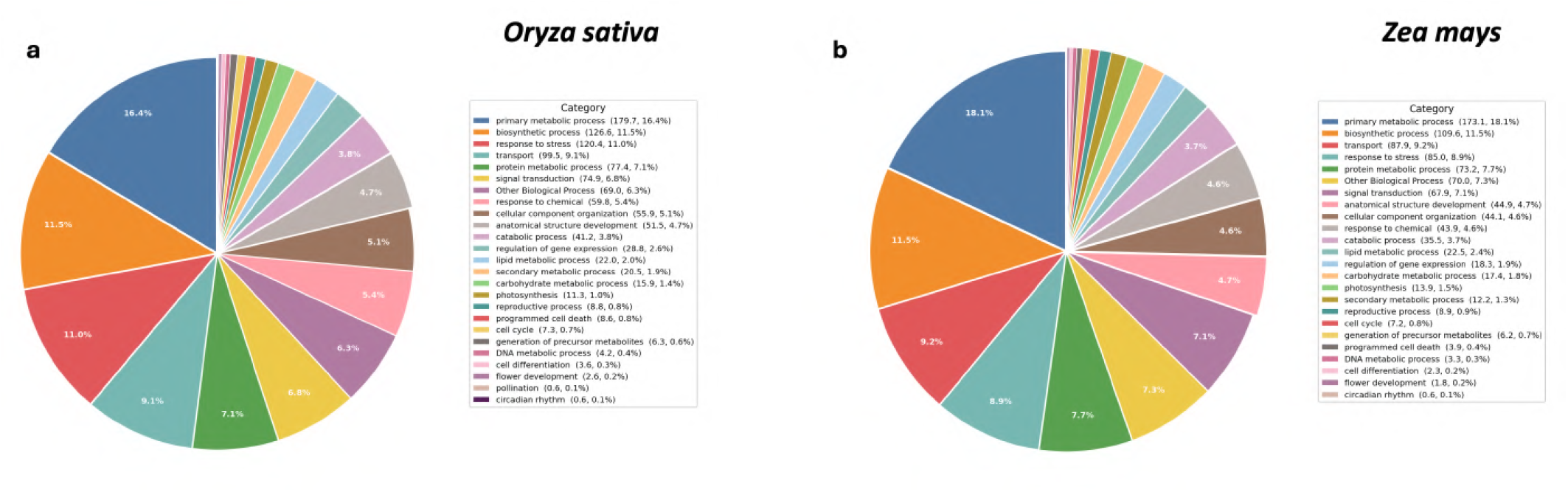
Gene Ontology analysis of predicted metamorphic proteins in representative plant proteomes. Distribution of high-level Gene Ontology (GO) biological process categories among predicted metamorphic proteins in (**a**) *Oryza sativa* and (**b**) *Zea mays*. GO terms were grouped into high-level biological process categories to facilitate comparison of the functional landscape of predicted metamorphic proteins across representative plant species.

**Fig. S10.**
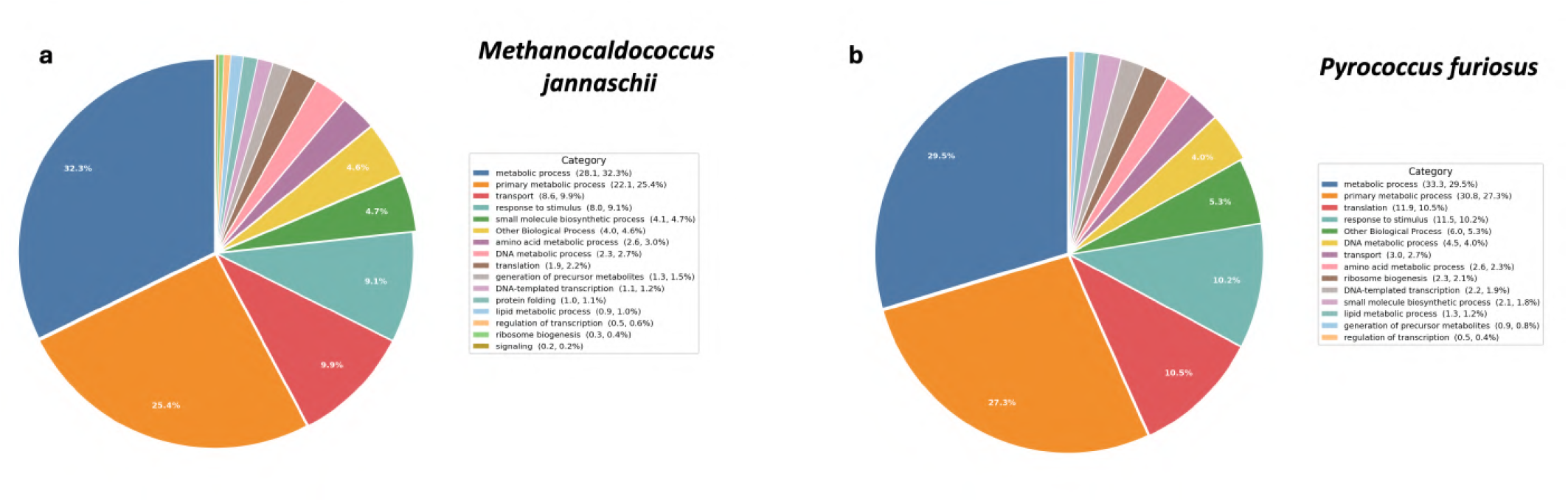
Gene Ontology analysis of predicted metamorphic proteins in representative archaeal proteomes. Distribution of high-level Gene Ontology (GO) biological process categories among predicted metamorphic proteins in (**a**) *Methanocaldococcus jannaschii* and (**b**) *Pyrococcus furiosus*. GO terms were grouped into high-level biological process categories to facilitate comparison of the functional landscape of predicted metamorphic proteins across representative archaeal species.

**Fig. S11.**
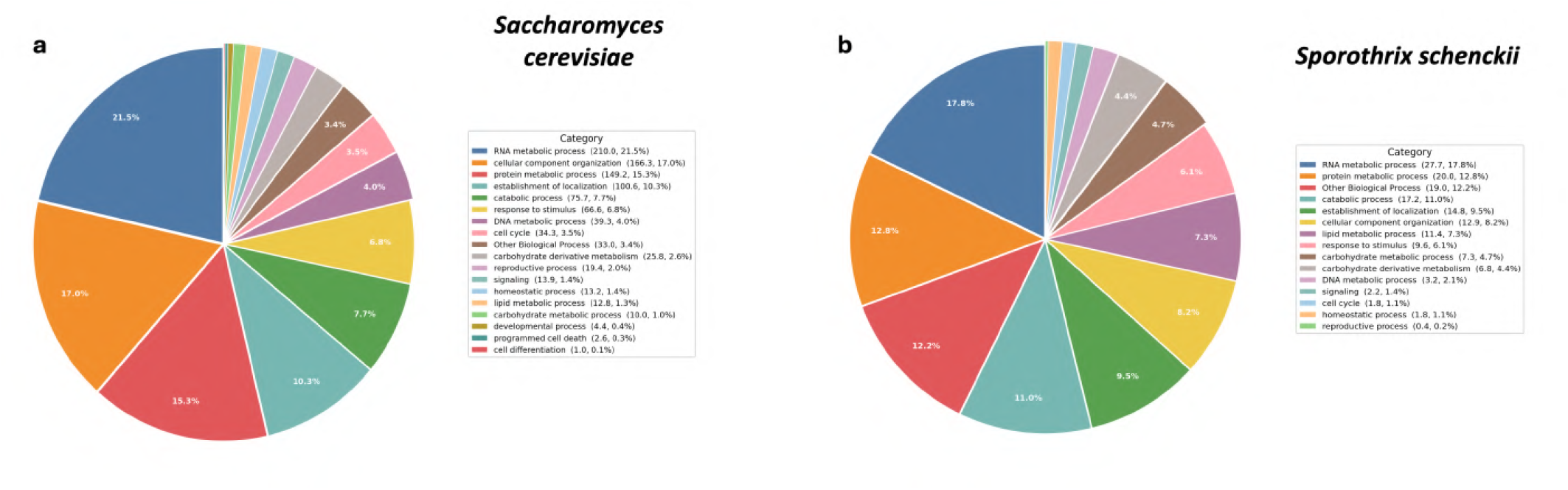
Gene Ontology analysis of predicted metamorphic proteins in representative fungal proteomes. Distribution of high-level Gene Ontology (GO) biological process categories among predicted metamorphic proteins in (**a**) *Saccharomyces cerevisiae* and (**b**) *Sporothrix schenckii*. GO terms were grouped into high-level biological process categories to facilitate comparison of the functional landscape of predicted metamorphic proteins across representative fungal species.

**Fig. S12.**
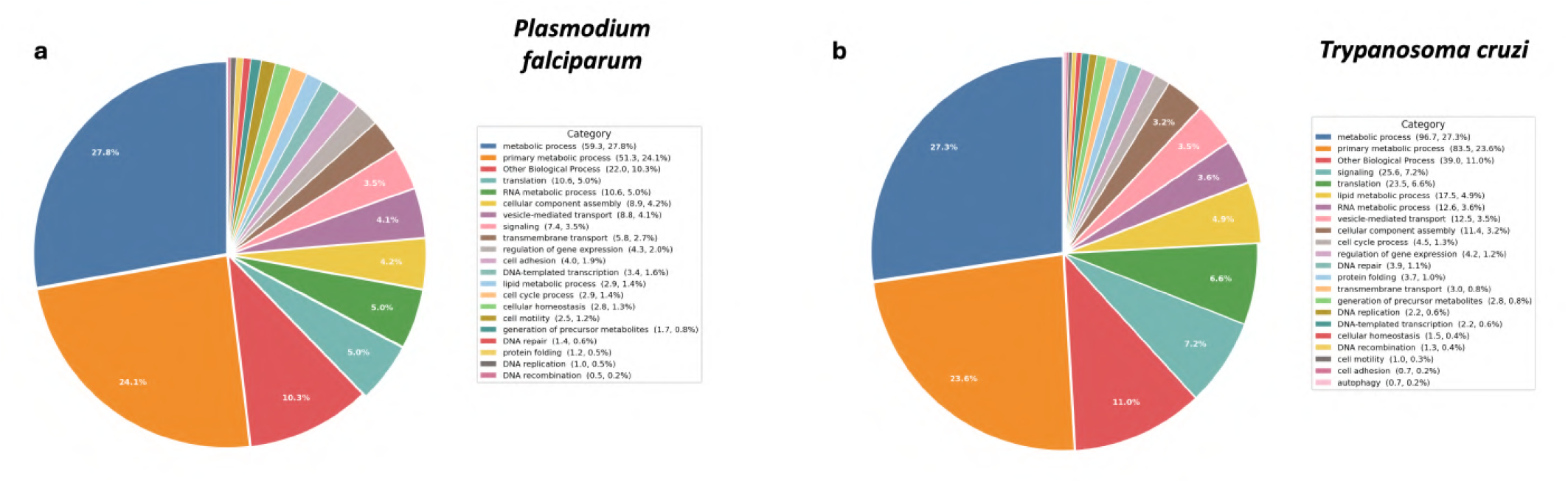
Gene Ontology analysis of predicted metamorphic proteins in representative protist proteomes. Distribution of high-level Gene Ontology (GO) biological process categories among predicted metamorphic proteins in (**a**) *Plasmodium falciparum* and (**b**) *Trypanosoma cruzi*. GO terms were grouped into high-level biological process categories to facilitate comparison of the functional landscape of predicted metamorphic proteins across representative protist species.

**Fig. S13.**
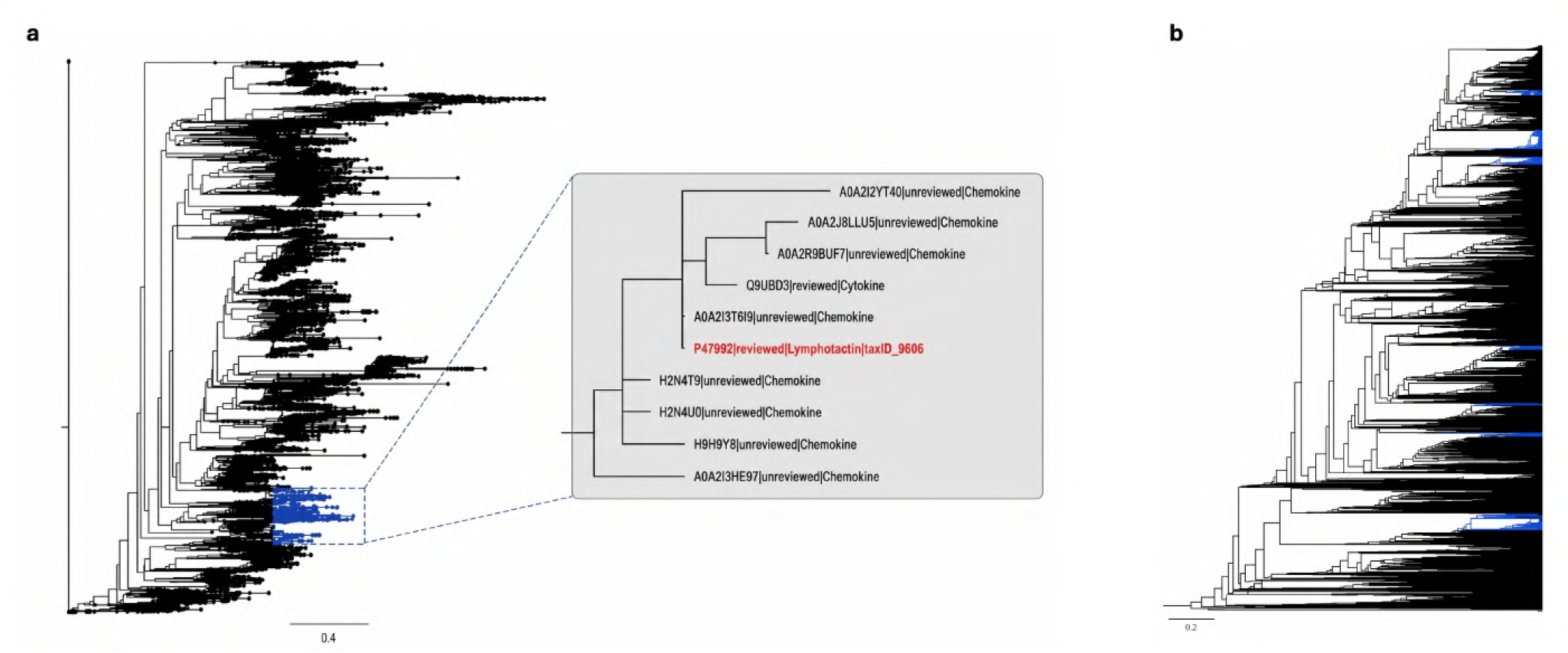
Evolutionary context of fold-switching protein families. (**a**) Maximum-likelihood phylogeny of the chemokine family showing the XCL1 (lymphotactin) lineage. The inset highlights the XCL1 clade and its closest homologues, illustrating the single evolutionary origin (monophyletic emergence) of the experimentally validated fold-switching chemokine. (**b**) Maximum-likelihood phylogeny of the RfaH/NusG superfamily. Branches containing proteins predicted as metamorphic by Morpheus-3D are highlighted in blue, revealing their distribution across multiple independent lineages, consistent with a polyphyletic origin of fold switching within the superfamily.

**Fig. S14.**
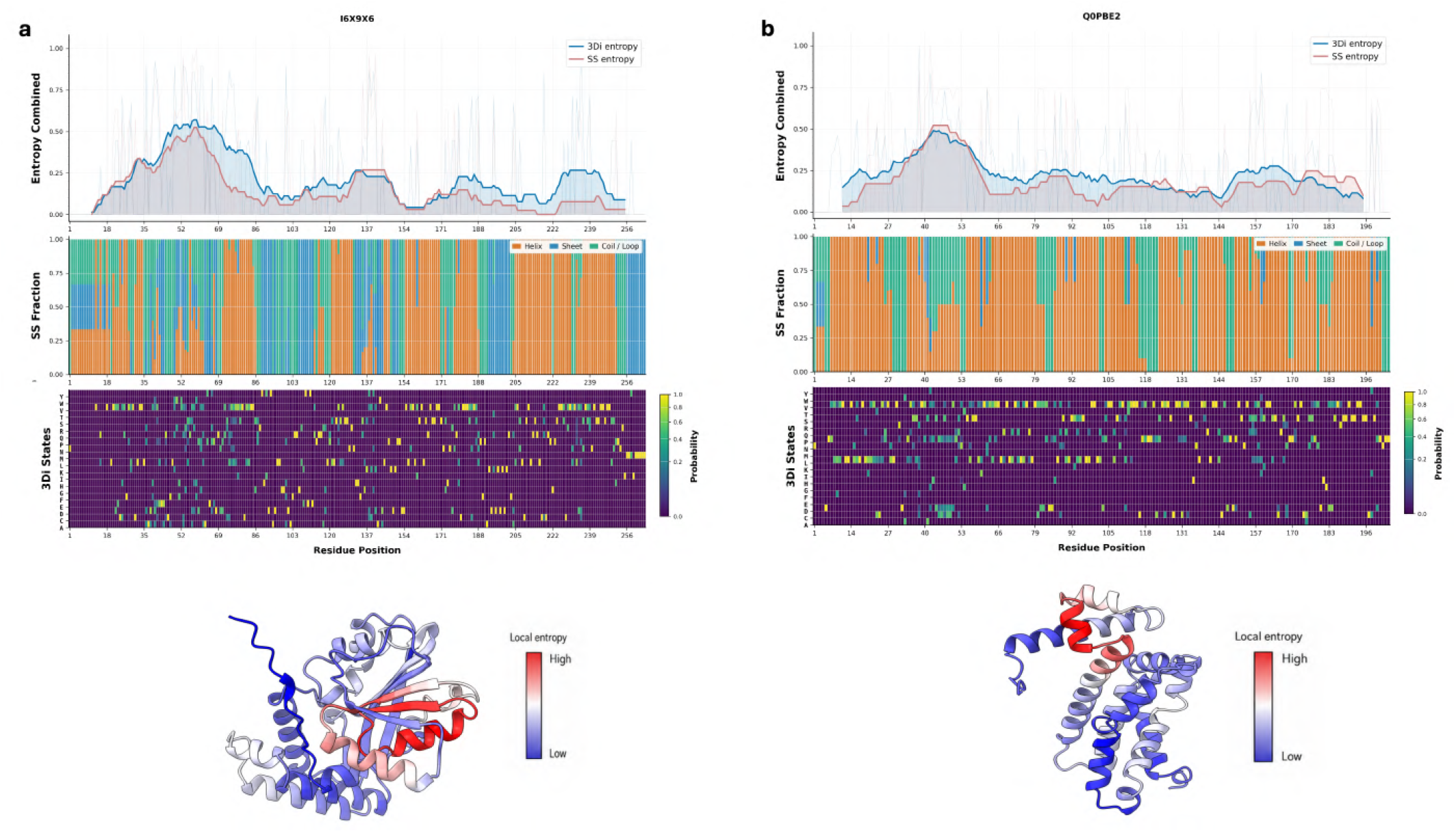
Structural diversity profiles of representative high-confidence metamorphic candidate proteins. (**a**) I6X9X6 (Methyltransferase domain-containing protein, *Mycobacterium tuberculosis*) and (**b**) Q0PBE2 (Transcriptional regulator CmeR, *Campylobacter jejuni*). The upper panels show rolling 3Di entropy (blue) and secondary-structure (SS; red) entropy profiles. Middle panels display the fractional occurrence of helix (orange), sheet (blue), and coil/loop (green) states together with residue-wise Foldseek 3Di state probabilities. The lower panels map normalized local 3Di entropy onto the predicted protein structures, where blue and red denote regions of low and high structural diversity, respectively, highlighting the predicted fold-switching regions identified by Morpheus-3D.

**Fig. S15.**
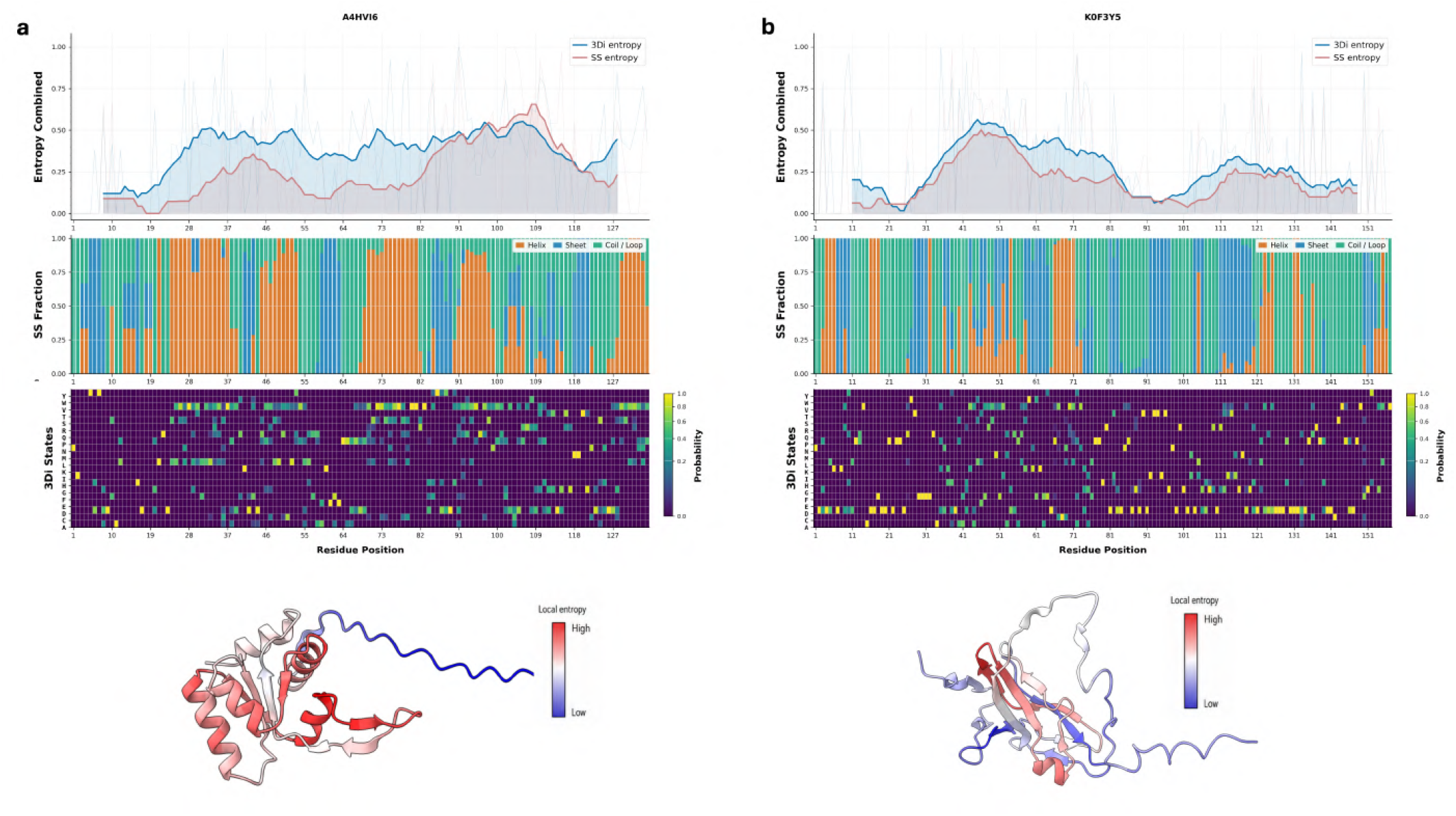
Structural diversity profiles of representative high-confidence metamorphic candidate proteins. (**a**) A4HVI6 (40S ribosomal protein S12, *Leishmania infantum*) and (**b**) K0F3Y5 (Deoxyuridine 5^*′*^-triphosphate nucleotidohydrolase, *Nocardia brasiliensis*). The upper panels show rolling 3Di entropy (blue) and secondary-structure (SS; red) entropy profiles. Middle panels display the fractional occurrence of helix (orange), sheet (blue), and coil/loop (green) states together with residue-wise Foldseek 3Di state probabilities. The lower panels map normalized local 3Di entropy onto the predicted protein structures, where blue and red denote regions of low and high structural diversity, respectively, highlighting the predicted fold-switching regions identified by Morpheus-3D.

**Fig. S16.**
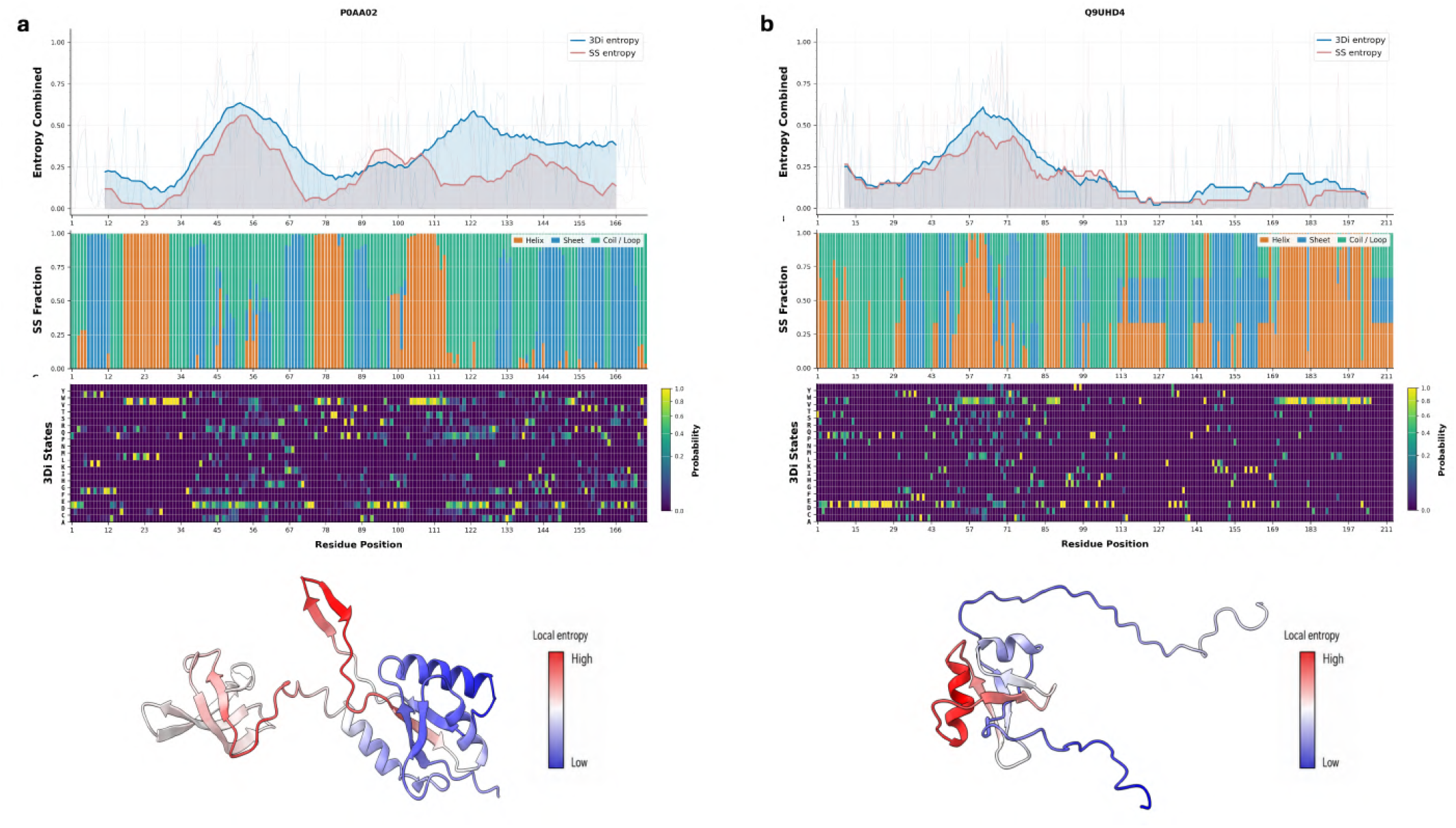
Structural diversity profiles of representative high-confidence metamorphic candidate proteins. (**a**) P0AA02 (Transcription termination/antitermination protein NusG, *Salmonella typhimurium*) and (**b**) Q9UHD4 (Lipid transferase CIDEB, *Homo sapiens*). Upper panels show rolling 3Di entropy (blue) and secondary-structure (SS; red) entropy profiles. Middle panels display the fractional occurrence of helix (orange), sheet (blue), and coil/loop (green) states together with residue-wise Foldseek 3Di state probabilities. Lower panels map normalized local 3Di entropy onto the predicted structures, where blue and red indicate regions of low and high structural diversity, respectively, highlighting the predicted fold-switching regions identified by Morpheus-3D.

**Fig. S17.**
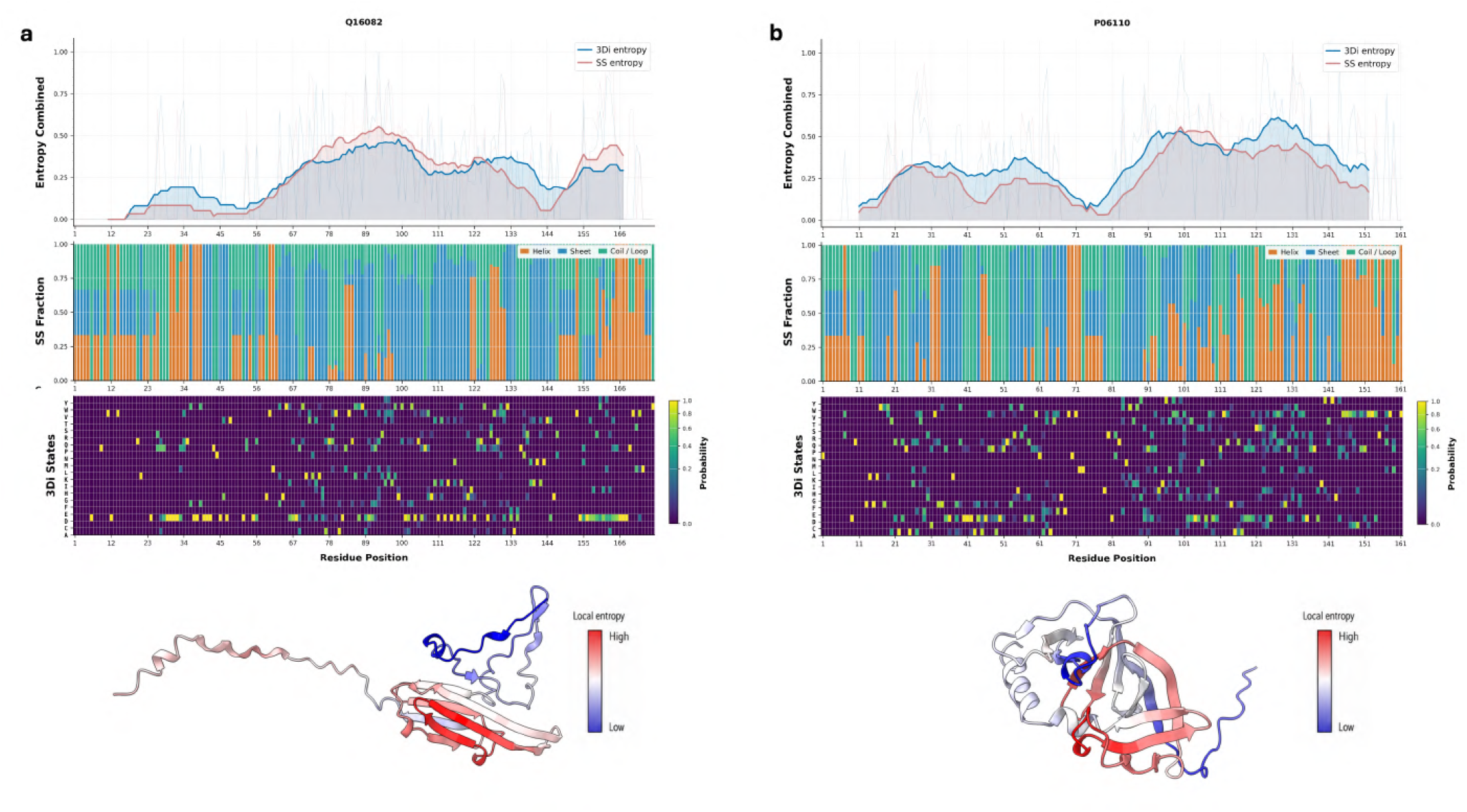
Structural diversity profiles of representative high-confidence metamorphic candidate proteins. (**a**) Q16082 (Heat shock protein beta-2 (HSPB2), *Homo sapiens*) and (**b**) P06110 (Chemotaxis protein CheW, *Salmonella typhimurium*). Upper panels show rolling 3Di entropy (blue) and secondary-structure (SS; red) entropy profiles. Middle panels display the fractional occurrence of helix (orange), sheet (blue), and coil/loop (green) states together with residue-wise Foldseek 3Di state probabilities. Lower panels map normalized local 3Di entropy onto the predicted structures, where blue and red denote regions of low and high structural diversity, respectively, highlighting the predicted fold-switching regions identified by Morpheus-3D.

**Fig. S18.**
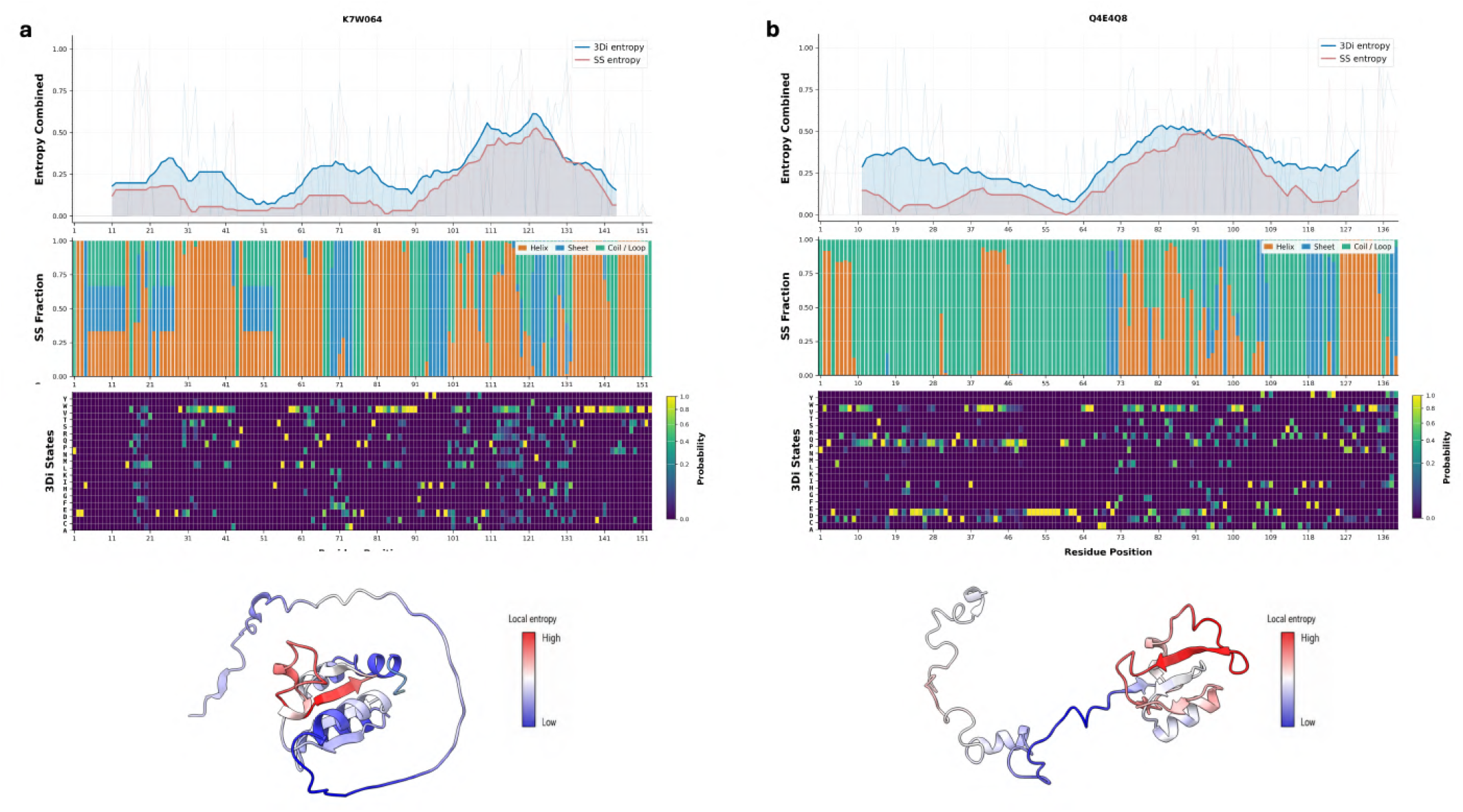
Structural diversity profiles of representative high-confidence metamorphic candidate proteins. (**a**) K7W064 (Glutaredoxin domain-containing protein, *Zea mays*) and (**b**) Q4E4Q8 (Large ribosomal subunit protein uL15, *Trypanosoma cruzi*). Upper panels show rolling 3Di entropy (blue) and secondary-structure (SS; red) entropy profiles. Middle panels display the fractional occurrence of helix (orange), sheet (blue), and coil/loop (green) states together with residue-wise Foldseek 3Di state probabilities. Lower panels map normalized local 3Di entropy onto the predicted structures, where blue and red denote regions of low and high structural diversity, respectively, highlighting the predicted fold-switching regions identified by Morpheus-3D.

**Fig. S19.**
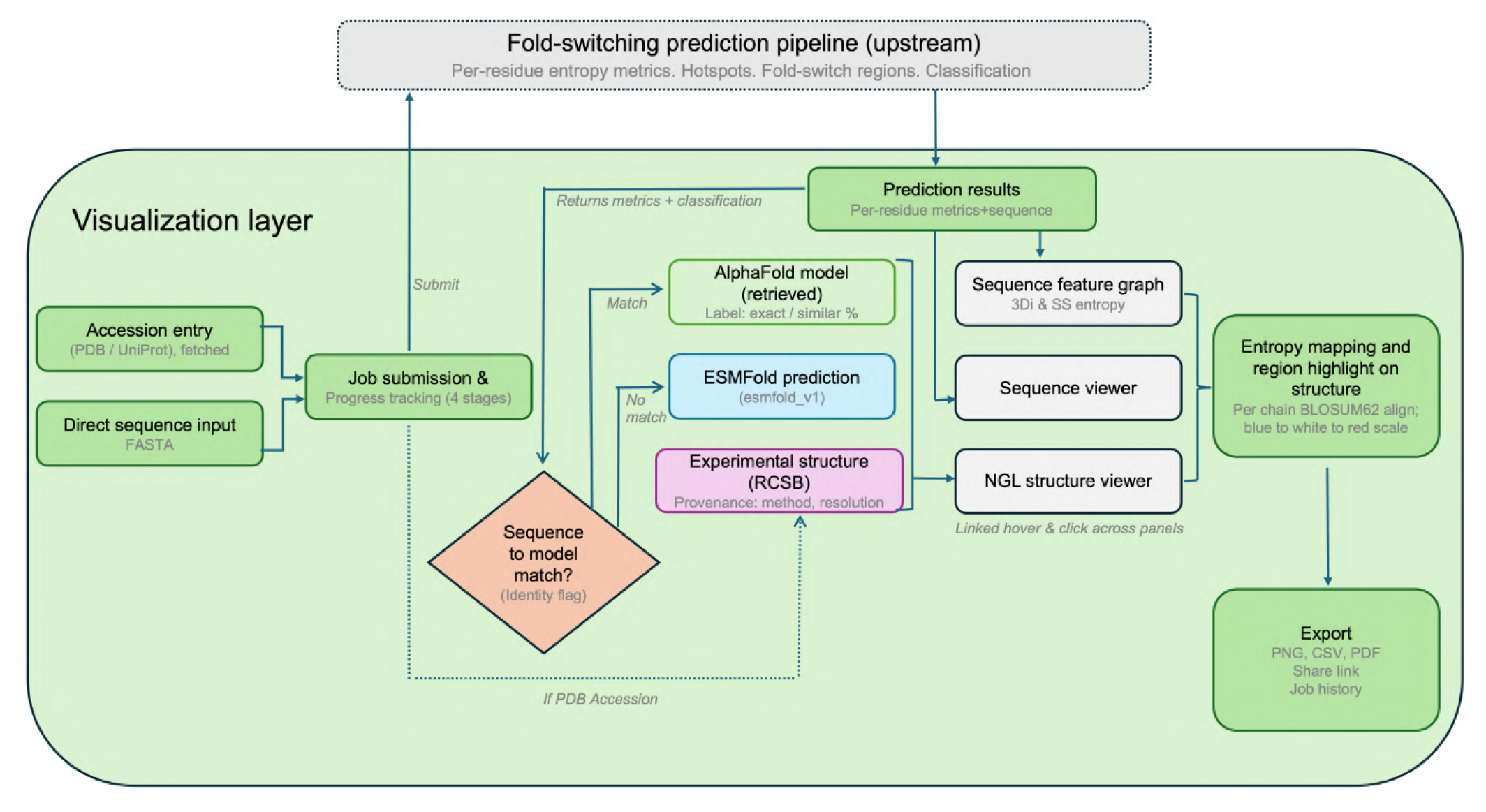
Workflow of the Morpheus-3D visualization layer. Morpheus-3D accepts a query as a direct amino acid sequence in FASTA format or as a PDB or UniProt accession. The submitted query is passed to the upstream fold-switching prediction pipeline, which returns per-residue entropy metrics, predicted hotspots, fold-switching regions and the overall classification; these results populate the sequence feature graph and sequence viewer. In parallel, the query sequence is tested against retrievable structural models to determine the structure source rendered in the NGL structure viewer: the corresponding AlphaFold-predicted structure is retrieved when an exact or highly similar sequence match is found, an ESMFold (esmfold_v1) structure is generated on demand when no match is found, and, for accession-based queries, the deposited experimental structure is additionally fetched from the PDB. The sequence feature graph, sequence viewer and structure viewer are linked through shared hover and click events, and the per-residue entropy profile is mapped onto the structure via a per-chain BLOSUM62 sequence alignment, coloured on a blue (low entropy) to white to red (high entropy) scale. Final outputs, including rendered structure views, per-residue metric tables and job history, can be exported as PNG, CSV or PDF, or retained as a shareable session link.

